# Ischemic injury triggers a protective microglial phenotype in models of Aβ pathology

**DOI:** 10.64898/2025.12.09.692939

**Authors:** Michael Candlish, Jan Hofmann, Desirée Brösamle, Annika Haessler, Murphy DeMeglio, Angelos Skodras, Georgi Tushev, Eloah S. De Biasi, Stefan Günther, René Wiegandt, Heidi Theis, Elena De Domenico, Nina Hermann, K. Peter R. Nilsson, Marc Beyer, Mario Looso, Maike Windbergs, Sigrun Roeber, Jochen Herms, Jonas J. Neher, Andreas G. Chiocchetti, Jasmin K. Hefendehl

**Affiliations:** Neurovascular Disorders, Institute of Cell Biology and Neuroscience, Biologicum, Goethe University Frankfurt, Max-von-Laue Str. 13, Frankfurt am Main, Germany; Department of Cellular Neurology, Hertie Institute for Clinical Brain Research, University of Tübingen, Tübingen, Germany; Biomedical Center (BMC), Biochemistry, Faculty of Medicine, LMU Munich, Munich, Germany; Neuroimmunology and Neurodegenerative Diseases, German Center for Neurodegenerative Diseases (DZNE), Munich, Germany; Munich Cluster for Systems Neurology (SyNergy), Munich, Germany; Institute of Pharmaceutical Technology, Goethe University Frankfurt, Max-von-Laue-Str. 9, Frankfurt am Main, Germany; Max Planck Institute for Brain Research, Max-von-Laue-Str. 4, Frankfurt am Main, Germany; Max Planck Institute for Heart and Lung Research, Member of the German Center for Lung Research (DZL), Member of the Cardio-Pulmonary Institute (CPI), Bad Nauheim, Germany; Platform for Single Cell Genomics and Epigenomics (PRECISE) at the German Center for Neurodegenerative Diseases and the University of Bonn, and West German Genome Center, Bonn, Germany; Immunogenomics & Neurodegeneration, German Center for Neurodegenerative Diseases (DZNE), Bonn, Germany; Department of Physics, Chemistry and Biology, Linköping University, SE-581 83, Linköping, Sweden; Center of Neuropathology and Prion Research, Faculty of Medicine, LMU Munich, Munich, Germany; Center for Neuropathology, Ludwig-Maximilians-University Munich, Munich, Germany; Department of Child and Adolescent Psychiatry, Psychosomatics and Psychotherapy, University Hospital, Goethe University Frankfurt Germany.

**Keywords:** Alzheimer’s disease, stroke, microglia, co-morbidity

## Abstract

Microglia are highly plastic cells that are capable of integrating subsequent insults. As the majority of Alzheimer’s Disease (AD) patients also show cerebrovascular pathology, we here aimed to dissect the interactions between AD and ischemic brain injury on the microglial response to amyloid beta (Aβ) pathology. Surprisingly, we find that ischemic stroke in the presence of cerebral β-amyloidosis results in the generation of a novel neuroprotective microglial phenotype. These microglia drive a rapid accumulation of highly dense Aβ plaques that exhibit a relatively benign nature and are strikingly similar to Aβ plaques observed in patients that are resilient to AD pathology. Thus, our data do not only highlight the impact of a co-morbid state of brain ischemia and Aβ pathology on the microglial phenotype but also identify novel molecular pathways that may serve to promote beneficial microglial functions in AD.

## Introduction

Alzheimer’s disease (AD) and cerebrovascular disease remain leading causes of mortality and morbidity. Notably, a significant proportion of AD cases exhibit signs of cerebrovascular pathology upon post-mortem examination^1^. Consequently, recent studies have aimed to investigate potential links between vascular dysfunction and subsequent cognitive impairment^2–4^, as it is essential to analyze the interaction of co-morbid disease states. Microglia, the resident immune cells of the brain, are critically involved in a range of neurodegenerative diseases^5,6^. They are pivotal in their response to stroke and AD but how they respond to the combined presence of both diseases remains uncertain, given their highly dynamic nature. Microglia possess the capacity to engage in both detrimental and reparative processes^7^ in the context of neurological disease depending on both the pathology and disease stage. In AD, along with other neurodegenerative diseases, microglia enter a disease-associated (DAM) phenotype^8^. In mouse models of cerebral β-amyloidosis, DAM cluster around amyloid beta (Aβ) plaques in AD, engage in Aβ phagocytosis^9^ and act as a physical barrier that restricts the expansion of the immature, neurotoxic Aβ halo surrounding mature (less neurotoxic) Aβ plaque cores^10^. As the pathology advances, there is an increased abundance of immature Aβ within the brain parenchyma, coupled with a decline in the formation of mature, dense-cored Aβ plaques and an increase of more toxic soluble prefibrillar Aβ^11^. This suggests a progressive decline in the protective function of microglia over time. Conversely, in stroke, microglia contribute to acute neuroinflammation, exacerbating neuronal loss, but also contribute to neuroprotection and post-ischemic repair. Importantly, the balance between these effects varies over the recovery period^12–14^. Consequently, microglia in ischemic stroke undergo temporally-regulated phenotypic alterations with opposing effects on the brain parenchyma in contrast to the apparent unidirectional phenotypic shifts seen in chronic neurodegenerative diseases. Taken together, the distinct pathological milieus of AD pathology and ischemic stroke in a co-morbid condition, hold the potential to modulate microglial response, with unknown influence on AD progression.

Here, we analyze the intricate interplay between ischemic stroke and AD pathology by utilizing the APPPS1 mouse model in conjunction with ischemic stroke. Using this murine co-morbidity model, we identify a microglia-dependent accumulation of Aβ plaques within the vulnerable peri-infarct region as rapidly as three weeks after stroke. This phenomenon occurred despite a well-established age/pathology-dependent cessation of *de novo* Aβ plaque formation^11^. Notably, newly-formed peri-infarct Aβ plaques exhibit a relatively benign nature, and are encapsulated by an overabundance of phagocytotic microglia. Using spatial and single cell RNA sequencing (scRNA-seq), we identified a dramatic expansion of two microglial clusters in the co-morbid model that were virtually absent in all other conditions. These clusters are characterised by increased phagocytic pathways and additionally, we identified a stark microglial upregulation of several S100 proteins known to promote Aβ aggregation. Taken together, our findings underscore that the pathological microenvironment created by ischemic stroke in AD triggers a functional switch in microglia that promotes the formation of highly compact Aβ plaques akin to those observed in patients that are resilient to the detrimental effects of AD pathology^15^

## Results

### Ischemic stroke triggers the accumulation of Aβ plaques proximal to the infarct border

To establish the impact of ischemic stroke on Aβ deposition and distribution within both the stroke core and surrounding tissue, we employed a photothrombotic stroke model in (5-18 months old) APPPS1 mice (a well-established model of Aβ deposition^16^). Aβ plaque distribution was subsequently analyzed at one, three- and nine-weeks post stroke (Fig. 1a). Additionally, we performed whole-brain tissue clearing using iDISCO at one- and three-weeks post stroke (Extended Data Video 1 and 2) in tandem with histological approaches. At one week post stroke we found no significant differences in mature dense-core (Methoxy_X04^+^) Aβ plaque load (percentage of brain area occupied by Aβ plaques) or the number of dense-core Aβ plaques in the peri-infarct tissue, suggesting no short-term effect on the examined pathological hallmarks (Fig. 1b-d, Extended Data Video 1). However, at three weeks post stroke, we observed a readily visible reduction in the number of Aβ plaques within the infarct core. In contrast, we detected a significant increase in both dense-core Aβ plaque load and number of dense-core Aβ plaques in the tissue adjacent to the infarct border (Fig. 1e-g, Extended Data Video 2). This phenomenon persisted up until nine weeks post stroke (Fig. 1h-j). Comparing the number of dense-core Aβ plaques within the 0-100 μm region around the infarct border at three weeks post stroke to animals with no stroke, we found that both the number of Aβ plaques and Aβ dense-core plaque load were significantly higher in the peri-infarct zone (Fig. 1k,l). To determine whether this phenomenon might be exclusive to the APPPS1 mouse model, we analyzed brains from a second model with cerebral β-amyloidosis, APP23 mice, three weeks after stroke (Extended Data Fig.1a). The APP23 mouse model features a higher ratio of Aβ40/42, considerably slower onset of Aβ deposition and kinetics as well as differences in Aβ plaque morphology compared to APPPS1 mice^16,17^. Strikingly, we found a similar accumulation of Aβ plaques in the peri-infarct region, suggesting that this phenomenon is independent of the model of cerebral β-amyloidosis (Extended Data Fig. 1b-d). It is important to note, that the formation of *de novo* dense-core Aβ plaques is virtually absent from approximately five months of age in APPPS1 mice without stroke (Extended Data Fig. 2a-m). Rather, there is a somewhat slow increase in the size of existing dense-core Aβ plaques^11^. To visualize immature Aβ species^18^, we used the luminescent conjugated oligothiophene (LCO) hFTAA (the emission spectra of which varies depending upon the Aβ structure to which they are bound^19,20^). Using this technique, we are able to infer structural information about Aβ conformation with spatial resolution. Using this dye, we identified an accumulation of toxic immature Aβ species (Extended Data Fig. 2a-m) over time. As all animals used in the present study were five months of age or older, our data demonstrate that ischemic stroke radically alters Aβ plaque accumulation and deposition kinetics within the vulnerable peri-infarct region as early as three weeks post stroke, resulting in the formation of new Aβ plaques and an apparently accelerated aggregation rate.

**Figure 1.**
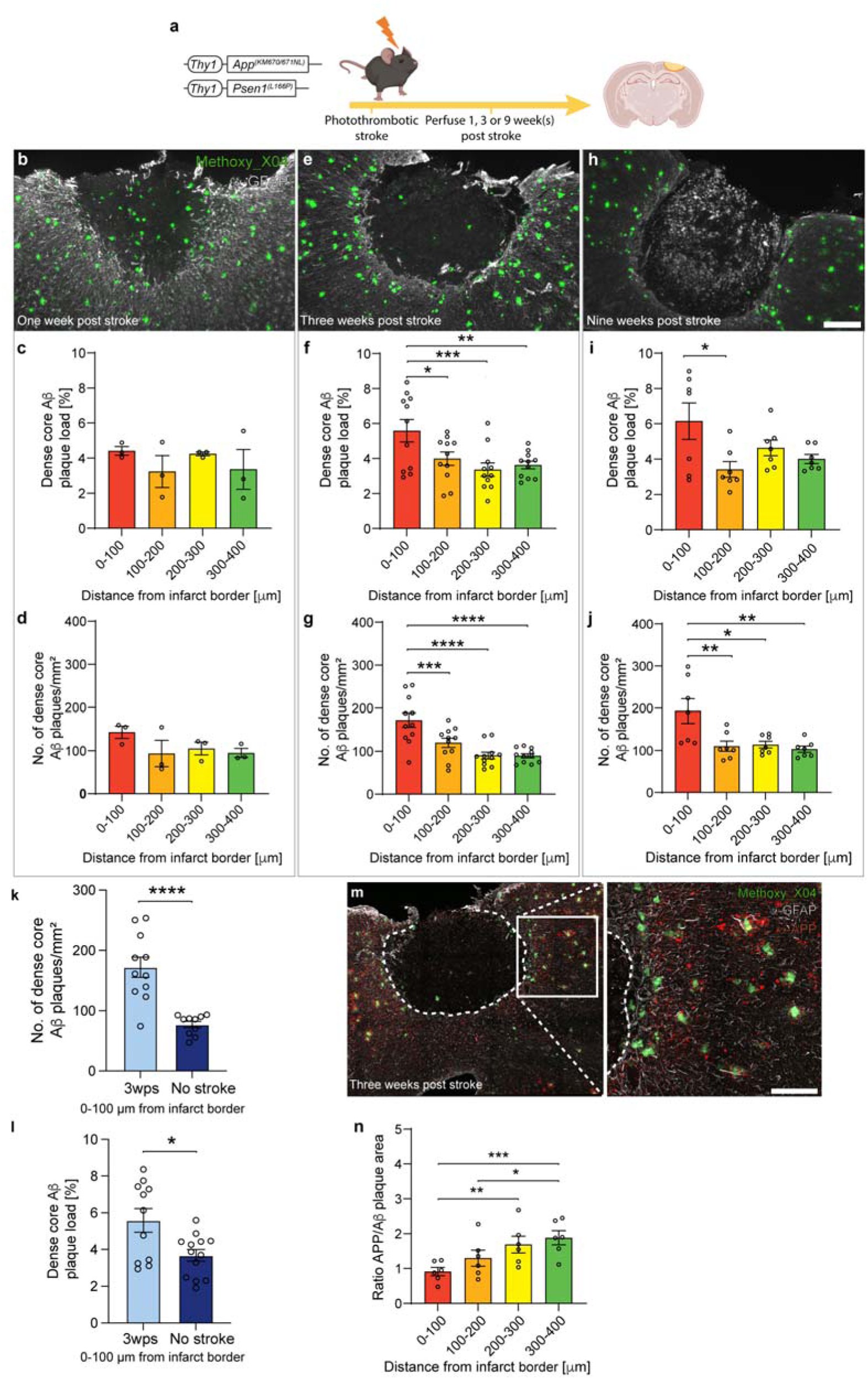
Ischemic stroke drives an accumulation of Aβ plaques in the peri-infarct zone. (**a**) Experimental strategy to model ischemic stroke in a mouse model of cerebral β-amyloidosis (APPPS1 mice). (**b**) Representative images of APPPS1 mouse brain sections one, (**e**) three and (**h**) nine weeks post stroke. Scale bar = 200 μm (n =3 mice; 1 male 2 females). (**c**) No significant difference in dense-core Aβ plaque load or (**d**) the number of dense core Aβ plaques in the peri-infarct region (n = 3 mice) at one week post stroke. (**f**) Significantly higher dense-core Aβ plaque load as well as (g) the number of dense core Aβ plaques were found in close proximity to the infarct border (n = 11 mice; 6 males 5 females) at three weeks post stroke. (**i**) Increased dense-core Aβ plaque load and (**j**) number of dense-core Aβ plaques persisted in close proximity to the infarct border up until at least nine weeks post stroke (n = 7 mice, 3 males 4 females). (**k**) The number of dense core Aβ plaques / mm^2^ (n = 11 three weeks post stroke mice, 6 males 5 females, n = 10, 5 males 5 females no stroke APPPS1 mice) and (**l**) the dense core Aβ plaque load is significantly higher 0-100 μm from the infarct border three weeks post stroke compared to APPPS1 mice without stroke. (n = 11 three weeks post stroke mice, 6 males 5 females, n = 13 no stroke APPPS1 mice, 7 males 6 females). (**m**) Representative image of an APPPS1 mouse brain section three weeks post stroke with APP immunolabelling to visualize dystrophic neurites. Scale bar = 50 μm (inset). (**n**) Significantly less Aβ-plaque associated axonal dystrophy was present in close proximity to the infarct border (n = 6 mice, 3 males 3 females). Dense-core Aβ plaques (b,e,h,m) are visible in green (labelled with Methoxy_X04), glial scar (b,e,h,m) is visible in white (GFAP immunoreactivity). Dystrophic neurites (**m**) are visible in red (amyloid precursor protein (APP) immunoreactivity). Repeated-measures one-way ANOVA with Tukey’s multiple comparison test for c, d, f, g, i, j and n. Unpaired two-tailed Student’s t-test for k,l. * = p < 0.05, ** = p <0.01, *** = p < 0.001, **** = p <0.0001.For full statistical details, see Supplementary Table 2.

### Peri-infarct Aβ plaques are structurally distinct and comparatively benign

We noticed that the peri-infarct Aβ plaques appeared morphologically distinct from the previously described Aβ plaques in APPPS1 mice, presenting with a more well-defined edge as opposed to a transition to immature neurotoxic Aβ halo as is typically seen around Aβ plaques in APPPS1 mice^21^. As more fibrillar (less compact) Aβ plaques have been linked to increased axonal dystrophy^10^, we hypothesized that peri-infarct Aβ plaques might display a lower neurotoxicity compared to “classical” Aβ plaques (non-stroke associated Aβ plaques). We therefore assessed the accumulation of the axonal dystrophy markers amyloid precursor protein (APP)^22^ as well as LAMP1 (Extended data Fig. 3a-b) in axons surrounding peri-infarct Aβ plaques. Indeed, we found significantly less dystrophic neurites surrounding Aβ plaques in close proximity to the infarct border from three weeks post stroke onwards (Fig. 1m,n, Extended Data Fig. 2n-s), suggesting that the peri-infarct Aβ plaques are comparatively benign.

To further interrogate the composition of the Aβ plaques in the peri-infarct area, we took advantage of the unique spectral properties of LCOs by using a combination of qFTAA and hFTAA, which have previously been utilized to resolve Aβ plaque maturation in AD mouse models^19,23^. By analyzing their emission spectra, structural information regarding the conformation of LCO-bound Aβ can be quantified^24^. To this end, we performed spectral scans on Aβ plaques labelled with qFTAA (which binds mature, compacted Aβ)^19,25^ and hFTAA (which binds immature/prefibrillar Aβ)^19,24^ one-, three-and nine-weeks post stroke (Fig. 2a-c). In the 0-400 µm region from the infarct border, we found that Aβ plaques were not only composed of more mature Aβ (as determined by the increased qFTAA to hFTAA ratio) as early as one-week post stroke but also surprisingly exhibited even greater compaction than Aβ plaques from APPPS1 mice without stroke (Fig. 2d-h). Accordingly, we also found a substantial reduction in the neurotoxic hFTAA positive Aβ halo surrounding Aβ plaques within the 0-100 µm perimeter around the infarct core starting at three weeks post-stroke and persisting until at least nine weeks post-stroke (Fig. 2i-l). Additionally, we observed a distinct loss of non-Aβ plaque associated small fibrillar Aβ deposits within the peri-infarct region (Fig. 2a-c). Given that Aβ can also be detected within neurons, we analyzed the intra to extra-neuronal distribution via surface reconstructions of the individual imaged components. We found that in APPPS1 mice, the vast majority of hFTAA-labeled Aβ is found extra-neuronally (Extended Data Fig. 3c-d).

**Figure 2.**
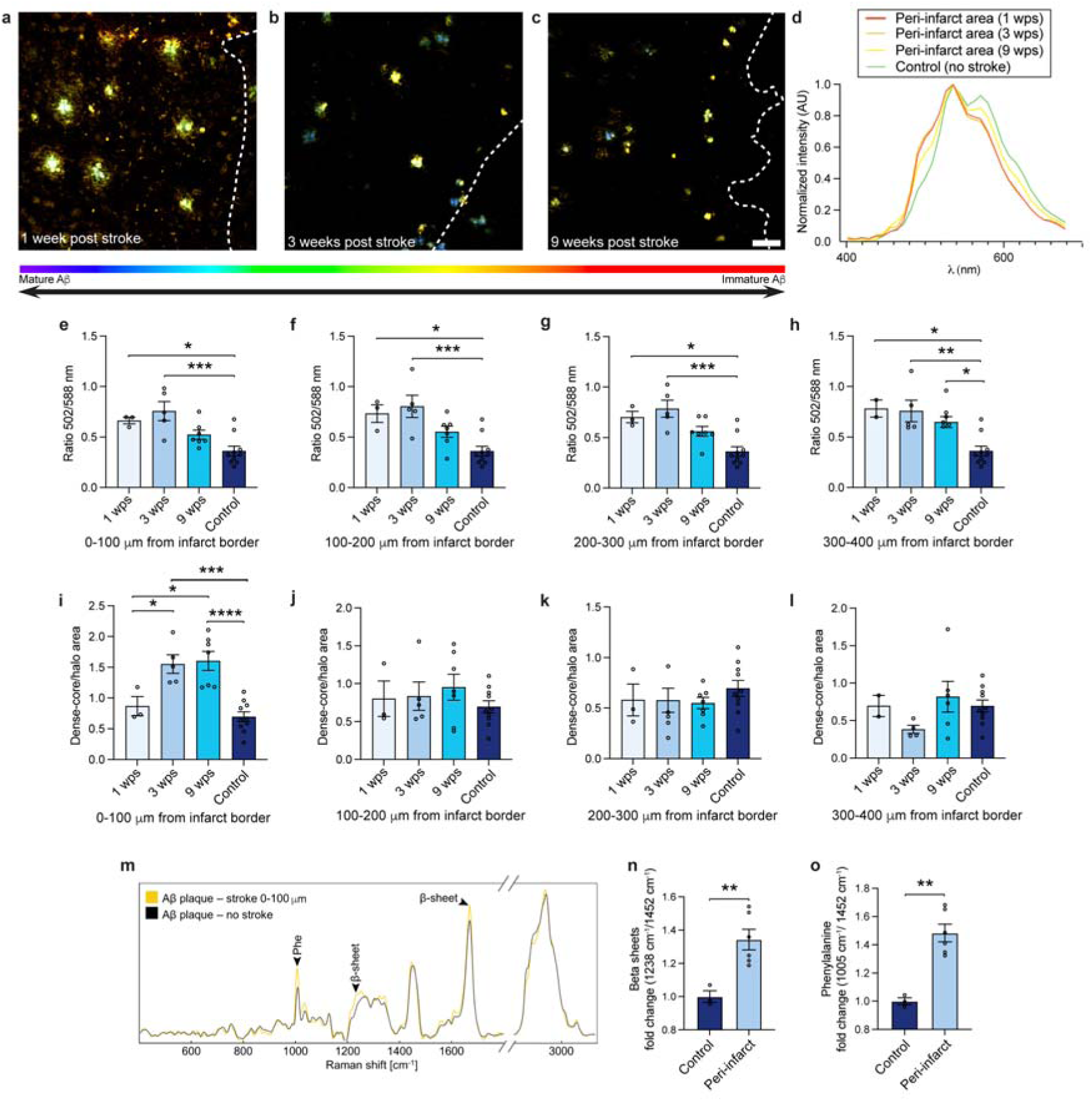
Peri-infarct Aβ plaques feature less neurotoxic immature Aβ halo and are more dense than typical Aβ plaques. (**a-c**) False-color coding of spectral scan images of APPPS1 brain sections stained with hFTAA and qFTAA taken at one (**a**), three (**b**) and nine (**c**) weeks post stroke (dashed lines demarcate the border of the infarct core, scale bar = 50 μm). Note the absence of immature Aβ halo as well as non-Aβ plaque associated immature Aβ deposits from three weeks post stroke onwards. (**d**) Representative normalized emission spectra traces of Aβ plaques in the peri-infarct area one-, three- and nine-weeks post stroke and without stroke. Note the general blue (i.e. left) shift of emission spectra in stroke compared to no stroke indicating increased density. (**e**) Dense core Aβ plaque to immature Aβ halo ratio was significantly increased compared to no stroke (and one-week post stroke) from three weeks post stroke onwards in the region 0-100 μm from the infarct border (**f,g,h**) but not distally (n = 2-10 mice; one week post stroke 2 male, 1 females, three weeks post stroke 3 males 2 females, 9 weeks post stroke 3 males, 4 females). (**i**) qFTAA/hFTAA spectral ratio at 502 nm and 588 nm (respectively) of peri-infarct Aβ plaques at one, three and nine weeks post stroke and with no stroke in APPPS1 mice (n = 2-9 mice; one week post stroke 2 males 1 female, three weeks post stroke 3 males 2 females, 9 weeks post stroke 3 males 4 females). A significantly higher qFTAA/hFTAA ratio was found consistently at one and three weeks post stroke compared to control (no stroke) at 0 – 100 μm, (**j**) 100 – 200 μm, (**k**) 200 – 300 μm and (**l**) 300 – 400 μm from the infarct border. (**m**) Averaged Raman spectra of peri-infarct Aβ plaques and Aβ plaques from APPPS1 mice without stroke. Markings indicate characteristic peaks of β-sheets and phenylalanine. (**n**) Peak ratio analysis of Raman spectra indicating β-sheets and (**o**) phenylalanine (n = 3 mice three weeks post stroke, two Aβ plaques per section (1 male 2 females), n = 3 mice no stroke APPPS1 controls, one Aβ plaque per section (2 males 1 female). Repeated-measures one-way ANOVA with Tukey’s multiple comparison test (e-l). Two-tailed unpaired t-test (j,k) * = p < 0.05, ** = p <0.01, *** = p < 0.001, **** = p <0.0001. For full statistical details, see Supplementary Table 2.

Next, to substantiate our findings we analyzed the compositional alterations in peri-infarct Aβ plaques using confocal Raman microscopy. As a label-free technique, it solely relies on the inelastic scattering of photons by molecular bonds, enabling the acquisition of spatially resolved specific molecular signatures. Consistent with our previous findings, we observed a significant increase in β-sheets as well as phenylalanine (a major component of the peptide primary and secondary structure, thus another indicator of density) within the peri-infarct Aβ plaques compared to typical Aβ plaques (Fig. 2m-o); confirming that ischemic stroke drives the generation of highly compact Aβ plaques in the peri-infarct region.

Our findings reveal dynamic changes in Aβ plaque pathology within the peri-infarct region induced by ischemic stroke, manifesting as heightened accumulation of mature Aβ plaques between one- and three-weeks post-stroke. This coincides with a discernible reduction in non-Aβ plaque associated neurotoxic immature Aβ species. These peri-infarct Aβ plaques exhibit a strikingly dense phenotype that exceeds the compaction of typical Aβ plaques. Furthermore, they feature a diminished fibrillar halo and crucially, induce less Aβ plaque-associated axonal dystrophy compared to typical Aβ plaques.

### Microglial encapsulation of peri-infarct Aβ plaques is augmented after ischemic stroke

To further study the underlying cause for the observed shifts in Aβ plaque deposition and aggregation kinetics, we next focused on microglia. They are of particular interest due to their putative role in Aβ plaque generation and compaction^20^, as well as their crucial role in restricting the spread of toxic immature Aβ halo^10^. In our co-morbidity model, we observed a notable increase not only in the Iba1^+^ area surrounding peri-infarct Aβ plaques near the infarct border (Extended Data Fig. 4a,b), but also a significantly increased microglia/Aβ plaque contact area on proximal peri-infarct Aβ plaques (Fig. 3a,b). Additionally, we found that the number of microglia surrounding peri-infarct Aβ plaques is higher proximal to the infarct core using PU.1 immunoreactivity to label microglial nuclei (Extended Data Fig. 4c,d). Furthermore, Aβ plaque-associated microglia proximal to the infarct border featured significantly higher CD68 intensity (a lysosomal marker expressed in phagocytotic microglia) (Fig. 3c,d). These observations suggest that microglia mount an augmented phagocytic response towards Aβ resulting from ischemic stroke. Note that the Iba1^+^ cells surrounding the peri-infarct Aβ plaques are TMEM119^+^, thus confirming their microglial identity^26^ (Fig. 3e).

**Figure 3.**
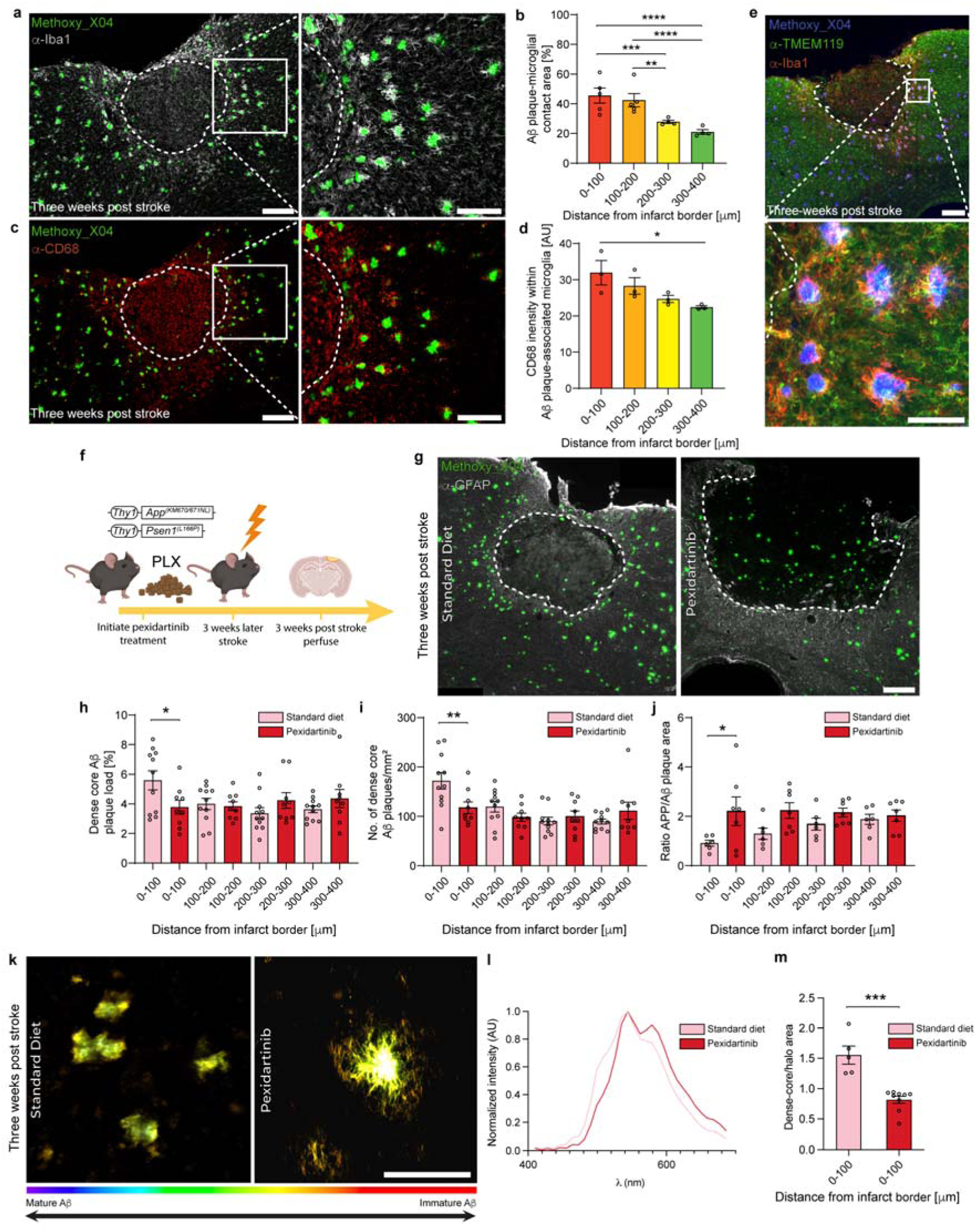
Microglia drive the formation of relatively benign Aβ plaques. (**a**) Representative image of an APPPS1 mouse brain section three-weeks post stroke with microglia (Iba1^+^ cells) visible in white and dense-core Aβ plaques (labelled with Methoxy_X04) in green. Scale bar = 200 μm, inset 100 μm. (**b**) Aβ plaque/microglial contact area (i.e. surface-to-surface contact area) is significantly higher in close proximity to the infarct border (n = 4-5 mice, 3 males 2 females). (**c**) Representative image of an APPPS1 mouse brain section three-weeks post stroke with the phagocytosis marker CD68^+^ cells visible in red and dense-core Aβ plaques (labelled with Methoxy_X04) in green. Scale bar = 200 μm, inset 100 μm. (**d**) Significantly higher CD68 intensity was detected in Aβ plaque-associated microglia in close proximity to the infarct border (n = 3 mice, 1 male 2 females). (**e**) Representative image of an APPPS1 mouse brain section three-weeks post stroke with microglia (Iba1^+^ cells) visible in red, the microglia-specific marker TMEM119 visible in green, and dense-core Aβ plaques (labelled with Methoxy_X04) in blue. Scale bar = 200 μm, inset 100 μm. (**f**) Experimental strategy to deplete microglia prior to ischemic stroke. (**g**) Representative image of an APPPS1 mouse brain section three-weeks post stroke after (right) microglial depletion with pexidartinib (PLX) and on standard diet (control). Note the lack of Aβ plaque accumulation in the peri-infarct area. Glial scar (GFAP immunoreactivity) is visible in white and dense-core Aβ plaques (labelled with Methoxy_X04) in green. Scale bar = 200 μm. (**h**) Pexidartinib treatment abolished the increase in dense-core Aβ plaque load and (**i**) the increased number of dense core Aβ plaques (n = 11 standard diet mice, 6 males 5 females, n = 9 pexidartinib treated mice, 5 males 4 females) (**j**) Pexidartinib treatment results in significantly more Aβ plaque-associated dystrophic neurites proximal to the infarct border (n = 6 standard diet mice, 3 males 3 females, n = 7 pexidartinib treated mice, 5 males 2 females). (**k**) False-color coding of spectral scan images of APPPS1 brain sections stained with hFTAA and qFTAA taken at three weeks post stroke following six weeks of pexidartinib treatment (dashed lines demarcate the border of the infarct core, scale bar = 50 μm). (**i**) Representative normalized emission spectra traces of Aβ plaques in the peri-infarct area at three weeks post stroke with and without pexidartinib treatment. Note the general red (i.e. right) shift of emission spectra resulting from pexidartinib treatment indicating decreased density. (**m**) Dense core Aβ plaque to immature Aβ halo ratio was significantly decreased after pexidartinib three weeks post stroke in the region 0-100 μm from the infarct border (n = 5 standard diet mice, 3 males 2 females, n = 9 pexidartinib treated mice, 4 females 5 males). Repeated-measures one-way ANOVA with Tukey’s multiple comparison test (b,c). Ordinary one-way ANOVA with Šidák’s multiple comparison test (h,j,j), two-tailed Students t-test (m). * = p < 0.05, ** = p <0.01, *** = p < 0.001, **** = p <0.0001. For full statistical details, see Supplementary Table 2.

To provide mechanistic evidence for the role of microglia in increased peri-infarct Aβ plaque formation, we next depleted microglia using the colony-stimulating factor 1 receptor (CSF1R) antagonist pexidartinib (Fig. 3f). Consistent with previous studies, we attained a significant, yet incomplete, reduction in microglia-occupied area since DAM^8^ no longer require CSF1R activation for their survival^20,20,27^ (Extended Data Fig. 4e-f). Remarkably, we found that pexidartinib treatment completely abolished increased peri-infarct Aβ plaque formation (Fig. 3g,h,i), indicating a dependence of this process on CSF1R-dependent microglia. Likewise, we found that peri-infarct Aβ plaques were less dense after microglial ablation and featured a lower dense-core Aβ plaque to halo ratio, reminiscent of typical Aβ plaques (Fig. 3k-m, Extended Data Fig. 4g). Crucially, axonal dystrophy was significantly higher around the Aβ plaques adjacent to the infarct core in pexidartinib-treated mice, further consolidating the importance of microglia in the described process (Fig. 3j, Extended Data Fig. 4h). Importantly, by analyzing axon density via neurofilament M labelling in brain sections from mice treated with either pexidartinib or standard diet, we found that there were no significant differences between both groups. These findings indicate that higher axonal dystrophy in pexidartinib treated mice is not a result of increased axonal survival after stroke compared to mice with an intact microglial population (Extended data Fig. 5a-b).

Using this depletion method, we show that CSF1R-dependent microglia play an essential role in the formation of inert peri-infarct Aβ plaques after stroke. It is worth noting however that the ablation of microglia may have indirect effects on Aβ plaque compaction by diminishing the clearance of myelin and cellular debris^28^.

### Co-morbidity-associated microglia (CoAM) are distinct from homeostatic- and disease associated microglia (DAM)

We next analyzed the molecular signature of the microglia driving the formation of relatively benign Aβ plaques in our co-morbid model, by analyzing their transcriptomic alterations. To this end we performed micro-dissections of brain tissue containing the infarct core along with the surrounding peri-infarct tissue and performed fluorescence-activated cell sorting (FACS) for microglia (excluding macrophages by gating for CD11b^high^ and CD45^low^ cells, as described previously^29^ (Extended Data Fig. 6a,b). To attain high quality transcriptomic data from the micro-dissected infarct and peri-infarct tissue, which limited the number of isolated cells, we used Smart-seq2 to delineate transcriptomic alterations in co-morbidity associated microglia (CoAM). Using this approach, we attained a highly enriched sort of microglia with only few non-microglial cells detected, which were subsequently removed prior to further analysis (Extended data Fig. 6c,d). Even though changes in expression levels were detected, all remaining cells used for subsequent analysis expressed typical microglial marker genes (i.e. *TMEM119, P2ry12, Cx3Cr1, Selplg*; Extended data Fig. 6e-j). We performed scRNA-seq (Fig. 4a) on microglia sorted from four different conditions: a) APPPS1 mice three weeks post-stroke (referred to as APPPS1 stroke), b) APPPS1 mice without stroke, c) WT mice three weeks post-stroke (referred to as WT stroke) and d) WT mice without stroke. When comparing microglia from all conditions using unsupervised clustering, we identified a total of seven clusters (Fig. 4b). In WT mice, as expected, the vast majority of microglia expressed homeostatic markers such as *P2ry12, Tmem119* and *Cx3cr1* (microglia clusters 2 (49.6%) and 4 (39.4%)) (Fig. 4a-c,f), as previously described^30^ (hence we refer to these clusters as homeostatic microglia 1 and 2, respectively). Conversely, in APPPS1 mice, the majority of microglia (48.2%) expressed genes such as *Tyrobp, Itgax, ApoE* and *Clec7a*, consistent with DAM (microglia cluster 0, hence we refer to this cluster as DAM) (Fig. 4a-c,f). In WT stroke, the most prominent microglial population (microglia cluster 1 (42.9%)) featured an upregulation of *Alpk1, Mmp12* and *Igf1*; genes previously identified to be upregulated by microglia in response to ischemia (Fig. 4a-c,f)^31–33^. Likewise, they also express some DAM genes such as *H2-D1*, *ApoE* and *Ctss*^8^ (Fig. 4f). Microglia cluster 1 together with microglia cluster 3 therefore, likely represent an intermediate stage between homeostatic microglia and DAM (hence we refer to cluster 1 and 3 as intermediate state microglia 1 and 2, respectively). Remarkably, and exclusively in the co-morbid condition, 46% of microglia formed microglia clusters 5 and 6 (25.1% and 20.9%, respectively) (Fig. 4a-c). In every other condition analyzed, these clusters did not exceed 8% of the microglial population (Fig. 4c), thus implicating these cells as co-morbidity associated microglia (CoAM). Both CoAM clusters featured upregulated *Dock10* and *Olfr334* whereas microglia cluster 5 almost exclusively expressed *S100b* (Fig. 4f). To attain an overview of the transcriptomic identity of these clusters, we split the cell populations into either homeostatic, DAM^+^/homeostatic^+^ (featuring markers of both homeostatic and DAM), DAM or neither signatures DAM^-^/homeostatic^-^, Fig. 4d,e). The vast majority of microglia in WT were enriched for the homeostatic signature (91.1%) with a small population determined to be neither DAM nor homeostatic (8.5%). As expected, in APPPS1 the majority of microglia showed a DAM signature (50.5%) and one third of microglia were classified as homeostatic. In WT stroke, there was an expansion of the DAM population (39.7%), but additionally an increase in microglia that were neither DAM nor homeostatic (22.2%) (Fig. 4d,e). However, in APPPS1 with stroke, the majority of microglial cells were in fact neither enriched for DAM or homeostatic signatures (for example *Apoe, Itgax, Tmem119, P2ry12*) (52.9%) (Fig. 4d,e). Furthermore, we analyzed whether the different clusters are enriched for known genetic subgroup marker signatures (e.g., interferon response microglia, axon tract microglia) but did not identify any matches. Consequently, a deeper characterization of this potential novel state was needed (Extended Data Fig. 7).

**Figure 4.**
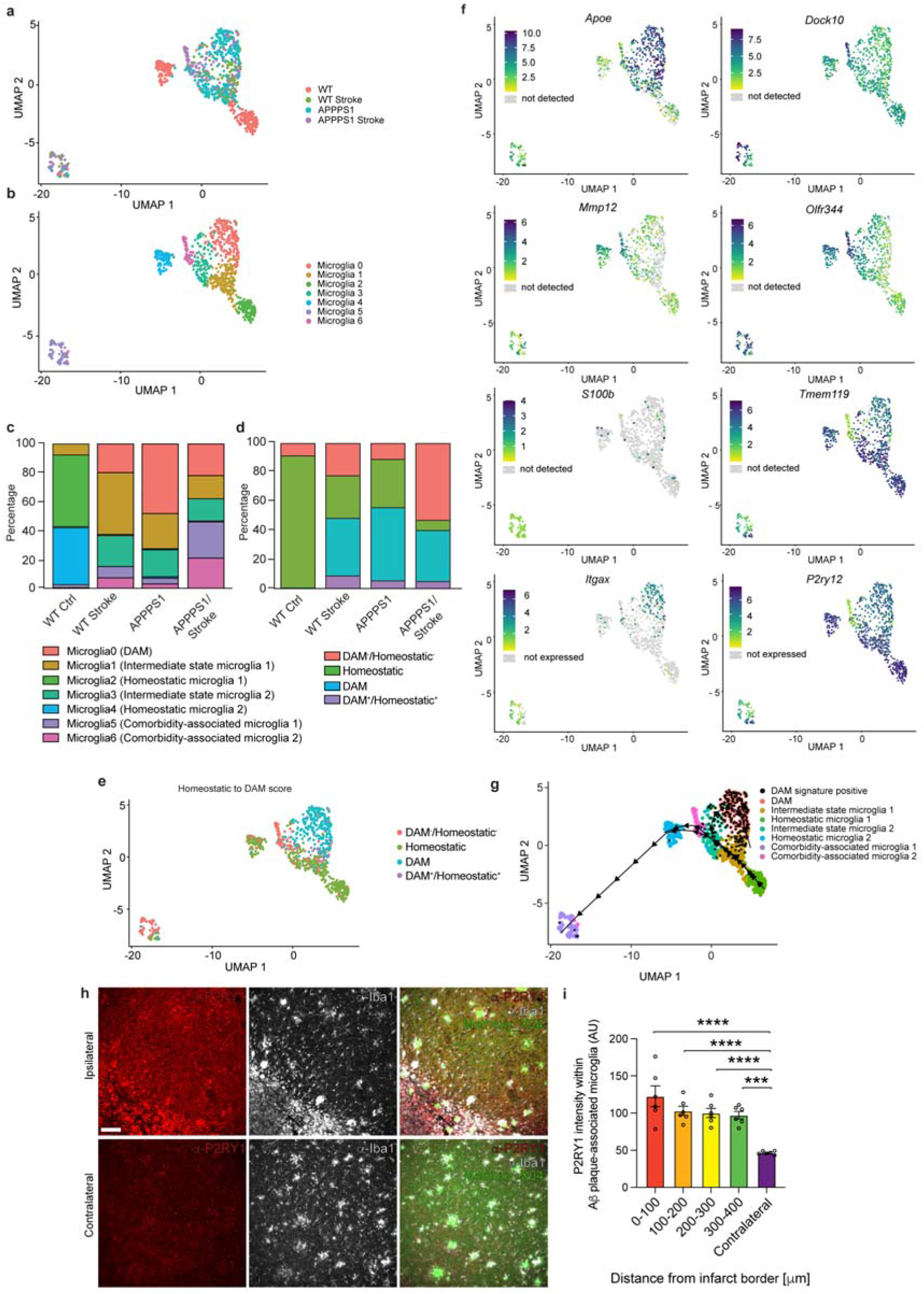
scRNA-seq reveals an over-representation of two microglial clusters in APPPS1 mice with stroke. (**a**) Uniform manifold approximation and projection (UMAP) plots of microglia from WT, WT stroke, APPPS1 and APPPS1 stroke mice. (**b**) UMAP plot of microglial clusters identified in WT, WT stroke, APPPS1 and APPPS1 stroke mice. (**c**) Stacked bar chart indicating the relative proportion of each microglial cluster in each condition. (**d**) Stacked bar chart indicating the relative proportion of microglia assigned as homeostatic, DAM, as having features of both DAM and homeostatic (DAM^+^/Homeostatic^+^) or being identified as neither DAM or homeostatic (DAM^-^/Homeostatic^-^). (**e**) UMAP plots of microglia indicating which microglia have been assigned to each microglia class as described in (d). (**f**) UMAP plots illustrating the expression levels of genes associated with homeostatic microglia (*Tmem119*, *P2ry12*), DAM (*Apoe*, *Itgax*) and genes found to be enriched in microglia that are neither DAM nor homeostatic (*Dock10*, *Mmp12*, *S100b*, *Olfr344*). (**g**) Trajectory analysis indicating transition from (1) homeostatic microglia 1 to DAM, (2) homeostatic microglia 1 to homeostatic microglia 2 and (3) homeostatic microglia 1 to CoAM. (**h**) Immunohistochemical labelling of P2RY1 (red), Iba1 (white) and methoxy_X04 (green) in close proximity to the infarct core (labelled ipsilateral) and on the contralateral hemisphere. (**i**) Quantification of P2RY1 intensity within microglia in close proximity to the infarct core and on the contralateral side. (n = 6 three weeks post stroke mice, 3 males 3 females). Scale bar = 80 μm. Repeated-measures one-way ANOVA with Šidák’s multiple comparison test. * = p < 0.05, ** = p <0.01, *** = p < 0.001, **** = p <0.0001. (For scRNAseq experiments, n = 3 wildtype mice (2 females, 1 male), n = 2 wildtype stroke (2 females), n=2 APPPS1 (2 females), APPPS1 stroke n = 2 (1 male, 1 female). For additional scRNAseq data, see Extended Data Table 1-10. For full statistical details, see Supplementary Table 2.

To delineate the transition from homeostatic microglia 1 to the different activation states, we used pseudotime analysis (Fig. 4g). The homeostatic cluster with the highest expression of typical markers for homeostasis (homeostatic microglia 1), was set as the point of origin in this analysis under the assumption that any stimulus-induced phenotype likely arises from these cells. We identified one trajectory from homeostatic microglia 1 to DAM (Fig. 4g) and, as expected, was characterized by the upregulation of hallmark DAM-associated genes, including *Cst7*, *Apoe* and *Clec7a* (Extended Data Fig. 8a). The second trajectory terminates within a CoAM cluster (microglia cluster 5) after transitioning through the other CoAM cluster (microglia cluster 6, Fig. 4g). Hence, we subsequently refer to microglia cluster 6 as CoAM 1 and microglia cluster 5 as CoAM 2. This trajectory is defined by alterations of genes associated with homeostasis such as *P2ry12*, *Ctss* and *Cst3* as well as changes to olfactory receptor genes such as *Olfr635* and *Olfr344* (Extended Data Fig. 8b). The third trajectory bridged between the two homeostatic microglial clusters, showing markers such as *Olfr635*, *Gnaz* and *Prickle2* (Fig. 4g, Extended Data Fig. 8c). Therefore, our data suggest that the transitions from homeostatic to either the DAM or CoAM clusters follow distinct trajectories. These findings are consistent with our data demonstrating that selective ablation of CSF1R-dependent microglia, but not DAM, with pexidartinib abolished the formation of highly dense peri-infarct Aβ plaques. Next, using the data attained from our scRNAseq, we next sought to visually identify CoAM within the peri-infarct region using immunohistochemical labelling of P2RY1, which was found to be enriched in CoAM compared to DAM. Consistent with our sequencing results we found that P2RY1 intensity was almost double in Aβ plaque-associated microglia within the peri-infarct region compared to those on the contralateral side (Fig. 4h-i), thereby confirming their identity on the protein level.

Diving deeper into the RNAseq data, we queried individual genes that are differentially regulated amongst all conditions (Fig. 5a) and additionally, that are differentially regulated in the CoAM clusters in comparison to DAM; since these transcriptional differences likely underlie the ability to generate highly compacted Aβ plaques (Extended Data Table 1). By comparing the identified CoAM clusters with DAM, it was apparent that both CoAM clusters are characterized by low expression of DAM-associated genes such as *ApoE*, *Axl*, *Tyrobp*, *Clec7a*, *Cst7* and *H2-D1* (Extended Data Table 1). We identified an upregulation of an abundance of *Olfr* genes; G-protein coupled receptors (GPCRs) that are expressed in olfactory sensory neurons but additionally play diverse roles in other tissues such as the brain^34^ and in diverse cell types including microglia^35^ (Extended Data Table 1). Notably, when comparing CoAM1 genetic signatures to all other clusters, several of these *Olfr* genes were identified as markers for CoAM 1 (Fig. 5b). Interestingly *Kcnq1ot1*, which has previously been shown to be upregulated in patients with transient ischemic attack^36^ and ischemic stroke^37^, was identified as a marker gene for CoAM 1. *Kcnq1ot1* is a long non-coding RNA crucially involved in autophagy^37^ – an important Aβ clearance mechanism in microglia^38^. While studies have suggested that enhanced autophagy may be detrimental in ischemic stroke alone ^37^, augmenting autophagy in AD has been proposed as a promising target to clear Aβ plaques^39^.

**Figure 5.**
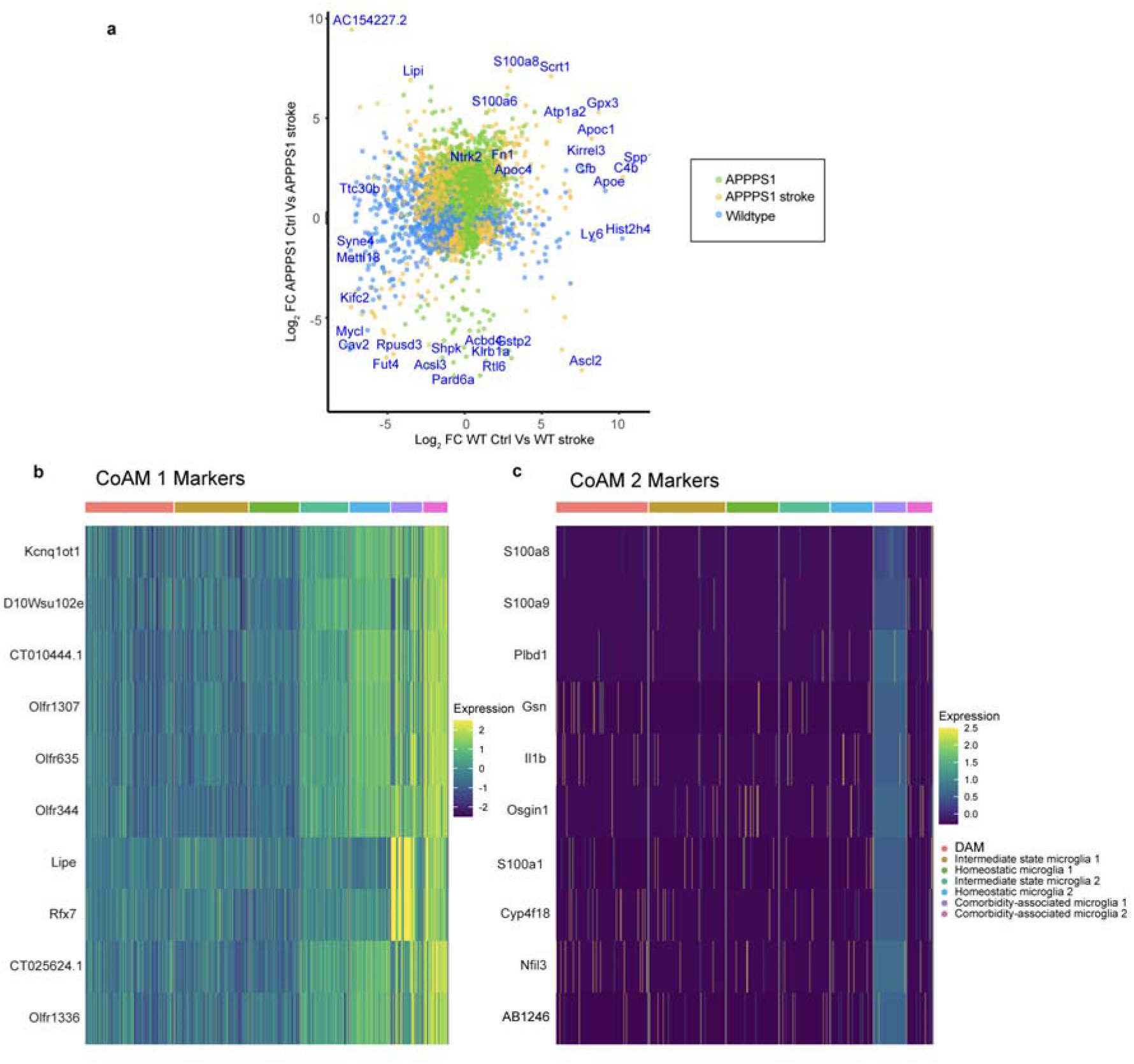
Differentially expressed genes associated with CoAMs. (**a**). Scatter plot showing differentially expressed genes in microglia from WT, APPPS1, WT stroke and APPPS1 mice. (**b**) Heatmap illustrating marker gene expression in CoAM 1 and (**c**) CoAM 2. Expression units = scaled log_2_ (counts +1).

Intriguingly, we also found a significant increase in leptin receptor (*Lepr*) expression, which has recently been implicated in microglial activation^40^ (Extended data table 1) in both CoAM 1 and CoAM 2. *Dock10* (which was amongst the most highly upregulated genes in CoAM 2 but also highly expressed in CoAM1, Fig. 4f) has been implicated in microglial cell migration^41^, potentially allowing CoAM to move to the sites of Aβ deposits. We further identified upregulation of *Sik3*; a gene involved in promoting microglial phagocytosis^42^ (Extended data table 1). Importantly, by comparing CoAM 2 to DAM, we found significant increases in many S100 genes (*S100a9*, *S100a8*, *S100a6*, *S100a10*, *S100a11*, *S100b*, *S100a7a*, *S100a3* and *S100pbp*; (Fig. 4f, Extended Data Table 1). Consistent with this, several S100 proteins were identified as marker genes for this cluster (Fig. 5c). S100 proteins have well-established, yet diverse and conflicting, roles in AD pathogenesis^43^. Crucially however, several of these proteins play a role in Aβ plaque aggregation^43^ and therefore, may be mechanistically involved in the formation of highly compacted Aβ plaques. Other marker genes we identified for CoAM 2 (Fig. 5c) include *Gsn* (involved in actin filament dynamics^44^ and therefore, potentially involved in microglial migration), *Il1b* (which has been implicated in augmented Aβ plaque removal^45^ and *Osgin1* (a transcriptional target of Nrf2, which has been demonstrated to be neuroprotective in a mouse model of AD) ^46^.

Leveraging the precise spatial localization of the ischemic stroke, we next employed spatial transcriptomics (Fig. 6a) on three-week post-stroke APPPS1 mice to validate the CoAM clusters we identified. Automated cell-type annotation indicated that invading macrophages were confined to the infarct core, while cells in the peri-infarct area were identified as microglia (Fig. 6b), confirming our previous results showing TMEM119^+^ microglia in the peri-infarct region (Fig. 3e) via immunostaining and identifying CoAM as microglia-derived. Moreover, clustering analysis demonstrated that spots containing microglia associated with distal Aβ plaques (blue) were clearly separable from spots containing peri-infarct Aβ plaque-associated microglia (orange) (Fig. 6c,d), the latter of which also demonstrated very tight clustering, indicating highly similar gene expression profiles (Fig. 6d). Comparing alterations between peri-infarct Aβ plaques and distal Aβ plaques (Fig. 6e, Extended Data Table 1), we identified several microglia-related genes that were upregulated in peri-infarct Aβ plaque-containing spots such as *ApoE*, complement proteins (*C1qa*, *C1qb*, *C3*) as well as lysosome-related genes (*Ctsb*, *Ctsd*). Additionally, we compared proximal and distal Aβ plaque-containing spots to spots identified as having a “homeostatic” microglial signature (green) in the APPPS1 model without stroke (i.e enriched for *P2ry12*, *Tmem119*, *Cx3cr1* and *Csf1r*). Consistent with our previous experiments demonstrating that the CSF1R-dependent microglial pool is the source of CoAM, we find that *Csf1r* is more abundantly expressed in peri-infarct Aβ plaque-containing spots than distal Aβ plaque-containing spots (Fig. 6e). Next, we compared differentially expressed genes between APPPS1 mice and APPPS1 stroke mice from our scRNA-seq dataset with genes upregulated in peri-infarct Aβ plaque-containing spots. As expected, we found a substantial number of mutually-expressed genes between the peri-infarct Aβ plaque-containing spots and APPPS1, since there are still DAM in this region (Fig. 6f). Additionally however, we identified 7 mutually expressed genes between APPPS1 stroke and peri-infarct Aβ plaque-containing spots (Fig. 6f). By examining the expression of these 7 genes in our scRNA-seq microglial clusters it became apparent that 100% of microglia in CoAM 2 expressed all 7 genes (Fig. 6g). Notably, CoAM 1 also expressed 4 out of 7 of these genes. Hence, the spatial analysis reinforces our earlier observations, that CoAM1 and CoAM2 are a unique microglial subtype with a remarkably beneficial phenotype that occurs as a result of the co-morbid state of cerebral ischemia and cerebral β-amyloidosis.

**Figure 6.**
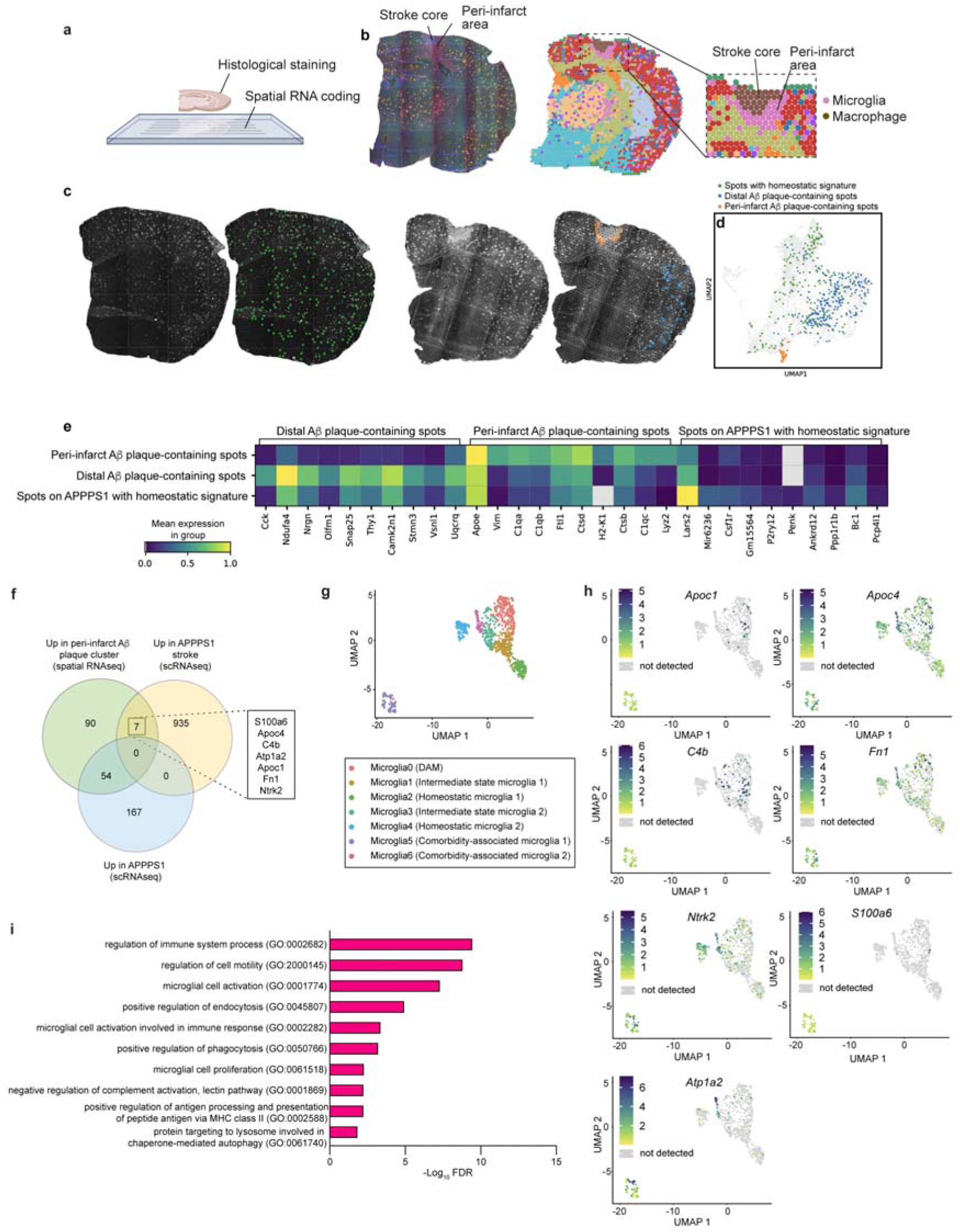
Spatial transcriptomics further implicates microglia clusters 5 and 6 as CoAM. (**a**) Graphical representation of spatial RNA-seq. (**b**) Automatic cell detection for microglia and macrophages. Note that macrophages are present within the infarct core whereas microglia are found throughout in the brain parenchyma including the peri-infarct area. (**c**) Immunohistochemical labelling of Aβ plaques (CN6), GFAP and microglia (CD45) on an APPPS1 coronal brain section from Visium spatial transcriptomics without stroke (left) and three weeks post-stroke (right) with representative images of automatically detected “homeostatic” microglia within a APPPS1 coronal section without stroke (left, shown in green) and manual selection of peri-infarct Aβ plaques (right, shown in orange) and distal Aβ plaques (shown in blue) for subsequent transcriptomic analysis. (**d**) UMAP showing clustering of all spots from n = 3 APPPS1 coronal sections three weeks post-stroke and one APPPS1 coronal section without stroke (from individual mice) used in spatial transcriptomic experiments. Homeostatic microglial signature-containing spots are shown in green, peri-infarct Aβ plaque-containing spots are shown in blue, distal Aβ plaque-containing spots are shown in orange. (**e**) Heatmap showing differentially expressed genes between peri-infarct Aβ plaque-containing spots, distal Aβ plaque-containing spots and homeostatic microglial signature-containing spots. (**f**) Venn diagram showing the overlap between differentially-expressed genes between APPPS1 stroke microglia (scRNA-seq) and APPPS1 microglia (scRNA-seq) and top upregulated genes in peri-infarct Aβ plaque-containing spots compared to distal Aβ plaque-containing spots. (**g**) UMAP plots of illustrating the expression levels of the seven mutually-upregulated genes between peri-infarct Aβ plaque-containing spots and APPPS1 stroke microglia (scRNA-seq). (**h**) Bar chart illustrating significantly enriched GO-terms in peri-infarct Aβ plaque-containing spots compared to distal Aβ plaque-containing spots. (For spatial transcriptomics, n = 6 three weeks post stroke mice, 2 females 1 male, n = 1 no stroke mouse, male).

By performing gene ontology (GO) analysis on the top upregulated genes in peri-infarct Aβ plaque-containing spots compared to distal spots, we found GO terms such as positive regulation of antigen processing and presentation of peptide antigen via MHC class II, negative regulation of complement activation (lectin pathway), microglial cell activation involved in immune response, protein targeting to lysosome involved in chaperone-mediated autophagy and positive regulation of endocytosis as well as positive regulation of phagocytosis to be highly enriched (Fig. 6h, Extended Data Table 1). These findings closely mirror pathways elucidated in a recent publication characterizing a microglial phenotype in patient samples exhibiting resilience to the toxic effects of Aβ deposition, in which a similar high compaction of Aβ is reported^15^.

In order to address the clinical relevance of our finding, we analyzed human brain sections to determine whether augmented Aβ plaque compaction also occurs in human’s post-stroke. To do this we analyzed human brain samples from three patients with ischemic stroke that also exhibited Aβ deposition. Despite the small sample number and an unclear timeline of the disease events, we found that human peri-infarct Aβ plaques were morphologically similar to the peri-infarct Aβ plaques identified in our mouse model (Fig. 7a-b). Interestingly, we observed a trend (below the threshold for statistical significance) towards reduced hFTAA volume (Fig. 7c) in the peri-infarct region, reflecting our findings in the mouse model. Consistent with this there was a trend towards increased qFTAA volume in the peri-infarct region in two out of three stroke patient samples (Fig. 7b,d), hinting towards a potential higher rate of compaction within the peri-infarct area. In an analysis of microglial engagement, we observed a trend towards increased microglial Aβ plaque coverage in the peri-infarct region. These human data support the hypothesis that the mechanisms uncovered in this study regarding improvement in microglia-mediated Aβ plaque compaction in mouse models may also occur in humans.

**Figure 7.**
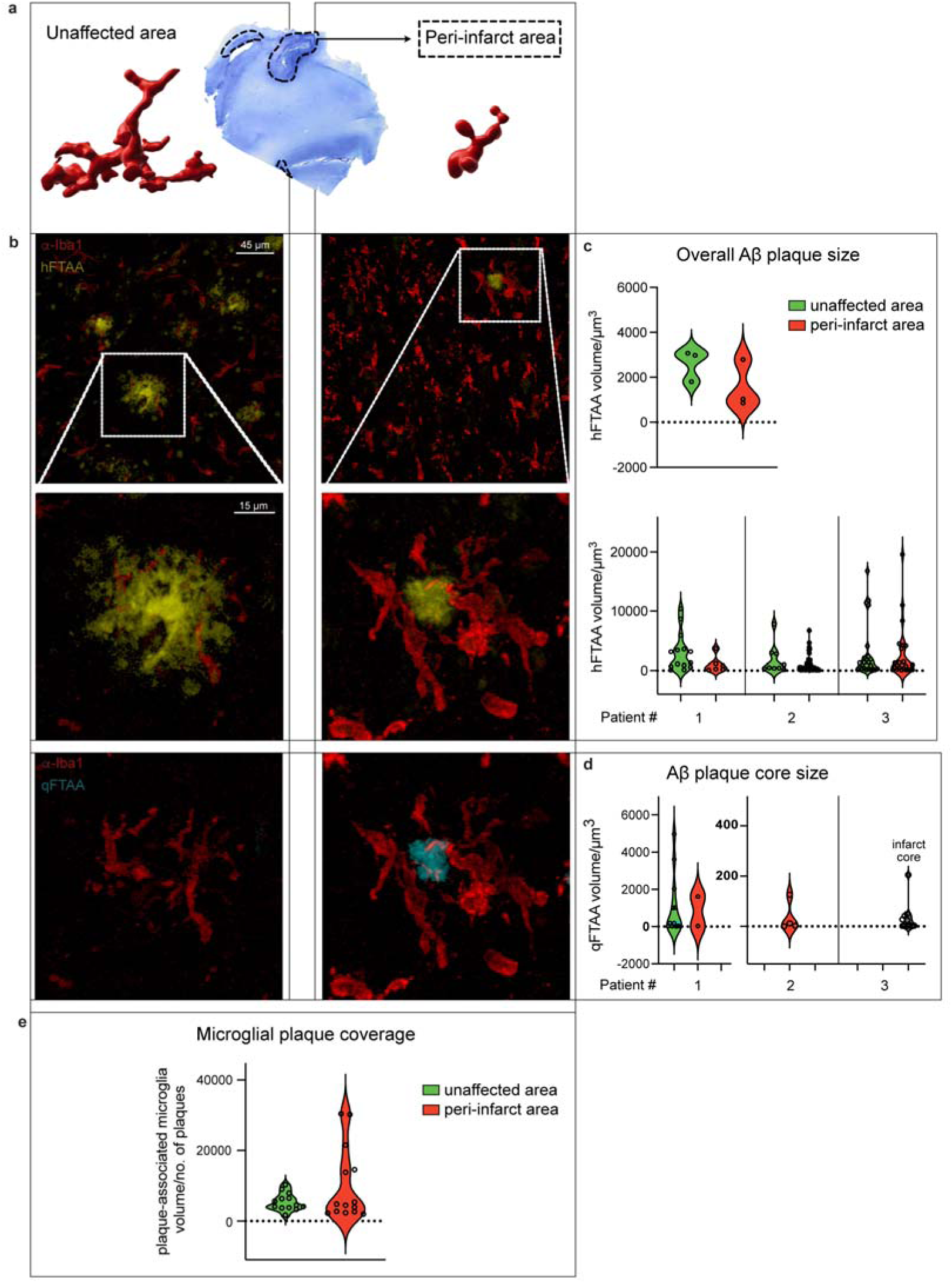
Peri-infarct Aβ plaques in humans are morphologically consistent with peri-infarct Aβ plaques in mice. (a) 3D reconstruction of microglia in human stroke tissue (Nissl stain shown in blue) and in unaffected brain regions. (b) Representative images of hFTAA- and qFTAA-labelled Aβ plaques in human stroke brain tissue. Note the striking morphological difference between Aβ plaques in the peri-infarct region compared to unaffected brain tissue. (c) Violin plot illustrating hFTAA volume of Aβ plaques in peri-infarct and unaffected tissue (top) with individual patient data shown below. (d) Violin plot illustrating qFTAA volume of Aβ plaque cores in individual patients. (e) Violin plot illustrating microglial Aβ plaque coverage in peri-infarct and unaffected brain tissue.

Taken together, these data indicate that the pathological milieu resulting from ischemic stroke and cerebral β-amyloidosis results in augmented microglial function, which drastically improves AD pathology by enabling microglia to generate comparatively inert Aβ plaques and to remove small fibrillar deposits of immature Aβ resulting in reduced axonal damage, reminiscent of the microglial-and Aβ plaque phenotype reported in resilient AD patients.

## Discussion

Modulation of microglial function is becoming an increasingly appealing target for disease-modifying treatment of neurodegenerative diseases^47^. Microglia can directly contribute to neuroinflammation in diverse conditions such as AD, Parkinson’s disease, multiple sclerosis, frontotemporal dementia, traumatic brain injury and ischemic stroke. Therefore, abrogation of neuroinflammation holds potential as a therapeutic strategy to prevent further neurological damage. Additionally however, microglia can also be beneficial in many of these conditions (such as via phagocytosis of cellular debris and protein aggregates such as Aβ in AD and α-synuclein in Parkinson’s disease for example). It is prudent therefore to investigate how microglia respond not only in each individual pathology but also in co-morbidity, since microglial function may be altered in the co-morbid state compared to a single pathology. Furthermore, many elderly patients in which these neurological diseases present have multiple comorbidities. Therefore, it is important to attain insight into the pathophysiological behavior of these cells both spatially – and temporally – as microglial phenotypes and cellular responses can vary not only between conditions but also over disease progression. By elucidating these idiosyncratic microglial phenotypes, we can develop a better comprehension of how we may be able to pharmacologically facilitate beneficial microglial phenotypes and to do this at the appropriate time in the course of each disease.

Investigating the effects of an ischemic insult, we here demonstrate that the somewhat dormant homeostatic microglial pool in cerebral β-amyloidosis model (and despite the abundance of DAM present in these mice), are able to regain beneficial functionality that allows them to phagocytically remove small fibrillar deposits and to compact large Aβ plaques; provided that they receive the appropriate external stimulus. It is important to note that the brain damage caused by ischemic injury certainly outweighs any potential benefit that could be attained from improved microglial function. Additionally, since the APPPS1 mouse model exhibits only mild cognitive impairment, a focal cortical stroke would be expected to dominate global behavior and mask any potential cognitive benefit resulting from augmented Aβ plaque compaction and reduced Aβ plaque-associated axonal dystrophy after stroke. Further work therefore must aim to pharmacologically promote this phenotypic alteration in microglial function in the absence of cerebral ischemia.

Within the infarct core we saw a nearly complete reduction in Aβ plaque load. Our data indicate that this phenomenon is mediated by macrophages confined to the infarct core. Thus, our focus remained on the accumulation of Aβ plaques in the peri-infarct region driven by brain-resident microglia. Strikingly, in addition to augmented dense-core Aβ plaque formation, we observed a stark loss of non-Aβ plaque associated deposits of immature/prefibrillar Aβ in the peri-infarct region. In APPPS1 mice (that develop Aβ plaques already at six weeks of age), new dense core Aβ plaque production is abrogated after five months of age followed by an accumulation of immature Aβ species in the brain parenchyma. Yet, despite this cessation of microglial function in terms of Aβ plaque compaction, the pathological milieu resulting from the co-morbid state of ischemic stroke with cerebral β-amyloidosis is sufficient to transform the remaining homeostatic microglia into a state of regained functionality, enabling them to create relatively benign dense-core Aβ plaques.

While the DAM state is initially useful to limit AD pathology, ultimately it is likely not beneficial as pro-inflammatory signaling gains the upper hand. This often has led to the characterization of the microglial response as a double-edged sword. However, we here demonstrate that this is not a unidirectional phenomenon for the entire microglial population in pathological settings; the remaining microglial population - when given the appropriate stimulus - is capable of undergoing a drastic phenotypic change leading to a functional improvement of the cells. Through our transcriptomic analysis we identified several potential mediators of this phenomenon. We identified a substantial increase in *Olfr* genes in CoAM. Recent studies have begun to investigate the role of olfactory receptors in microglial physiology. *Olfr110*, for example, has been found to be upregulated in models of neurodegenerative disease^48^. Likewise, pathogen-derived metabolites have been found to activate microglia via olfactory receptors^35^. It is important to note however, that our understanding of microglial olfactory signaling remains in its infancy and demands further exploration. Intriguingly, we identified an upregulation of the receptor for leptin, a hormone primarily secreted by white adipose tissue to control food intake. Leptin administration has recently been described to drive microglial activation, increase Iba1^+^ cell number^40^ and even elicit a neuroprotective effect in APPPS1 mice^49^. This phenotype is consistent with our findings. Additionally, we identified a clear upregulation of S100 proteins that have a well-established role in Aβ plaque formation. The roles of these pro-inflammatory proteins in neurodegenerative diseases are under current investigation, yet it remains unclear whether the expression of particular S100 proteins (or their pattern and expression level) are beneficial or not. For example, S100A11 has been demonstrated to be neuroprotective and improve cognition after ischemic stroke^50^. S100b overexpression results in increased cerebral β-amyloidosis^51^ but at lower concentrations inhibits Aβ plaque aggregation via interactions with Aβ_42_ monomers, oligomers and fibrils with a concomitant protection against cytotoxicity^52^. Likewise, S100A9 has been shown to interact with Aβ to induce fibrilization and is expressed in CD68 positive microglia after traumatic brain injury^53^. Also, several of these proteins (S100B, S100A1, S100A8 and S100A12) have been found in Aβ plaques^43^. Based on our results here – considering that we found reduced axonal dystrophy associated with peri-infarct Aβ plaques – it is tempting to speculate that augmenting Aβ plaque formation and compaction may be mediated via increased S100 protein expression. This might make it possible for Aβ plaques to act as a more efficient sink for toxic Aβ species, thus attenuating Aβ-associated neurotoxicity.

Most importantly, this study opens up a clear opportunity for future translational research. This is further reinforced by our findings in human stroke patients that also exhibited amyloid deposition. Given that the timeline of disease events is unclear in the analyzed patients a high variability in the measured parameters was to be expected. Nonetheless, our findings suggests a higher microglia coverage of Aβ-deposits in the peri-infarct area enabling them to form highly compact Aβ plaques. This could be seen by a trend in decreased hFTAA labeling and an increase in qFTAA labelling – both indicating an accelerated compaction rate within the analyzed area. Additionally, a recent publication documented an overabundance of highly compact Aβ plaques in AD patients that are seemingly resilient to Aβ toxicity and are therefore described as asymptomatic^15^, similar to what we found in our co-morbidity model. Based on these observations it is possible that the formation of highly compact Aβ plaques may be the biological basis of asymptomatic AD. If this is indeed the case, our findings suggest that microglia could be targeted pharmacologically to promote the transcriptomic and phenotypic changes we observed in our co-morbidity model, thus rendering Aβ plaques benign and preventing cognitive decline.

## Supporting information

Extended Data Table 1 - DEGs and GO term analyses

Extended Data Table 2-Pseudotime1_genes

Extended Data Table 3-Pseudotime2_genes

Extended Data Table 4-Pseudotime3_genes

Extended Data Table 5-DEG_Genelist_Microglia2vsMicroglia4

Extended Data Table 6-DEG_Genelist_Microglia5vsMicroglia6

Extended Data Table 7-DEG_Genelist_Microglia3vsMicroglia2

Extended Data Table 8-DEG_Genelist_Microglia5vsMicroglia0

Extended Data Table 9-DEG_Genelist_Microglia6vsMicroglia0

Extended Data Table 10-Clusters_Markers_all

Supplementary Table 1 - Antibodies

Supplementary Table 2 - Statistics

## Author contributions

M.C, J.H, and J.K.H conceived the study. M.C, J.H, A.H, M.D, G.T, E.S.D.B, D.B, A.S, A.G.C, R.W, H.T, E.D, N.H and S.G conducted experiments and/or analyzed data. S.R, J.H and K.P.R.N contributed essential reagents. M.L, M.B, M.W, A.G.C, S.R, J.H. K.P.R.N and J.J.N contributed expertise and feedback. M.C, J.H, and J.K.H wrote the manuscript.

## Acknowledgements

We thank Prof. Dr. Amparo Acker-Palmer for resource sharing and support of this project, Dr. Paolo Giacobini for technical advice regarding the iDISCO protocol, Almas Mahmood, Gisa Prange and Elif Ertas for excellent technical assistance, Guido Schmalbach for preparing the 3D printed iDISCO slides, and Lucas Ramirez Posada and Konstantinos Stefanidis for their work on the iDISCO experiments as part of their MSc rotation work. J.J.N. received funding from the Deutsche Forschungsgemeinschaft (DFG, German Research Foundation), Munich Cluster for Systems Neurology (EXC 2145 SyNergy – ID 390857198). J.K.H received funding from the Deutsche Forschungsgemeinschaft (DFG, German Research Foundation)-4b03813475, 419157387, Emmy Noether Award (HE 6867/3-1), SFB 1531 –Projektnummer 456687919, the Alzheimer’s Association (AARF-17-529810), and the Alzheimer Forschung Initiative e.V. (20041). BMBF (FK:01EW2308A) Neuron-ERANET, Cardio-Pulmonary Institute (CPI), EXC 2026, Project ID: 390649896 and CRC1080 to Prof. Dr. Amparo Acker-Palmer (221828878).

## Data availability

Any additional requests for data should be addressed to Prof. Jasmin Hefendehl.

## Code availability

The code for segmentation of qFTAA- and hFTAA-labelled Aβ plaques can be found here: https://gitlab.mpcdf.mpg.de/mpibr/scic/roioverstack. Any additional requests for code should be addressed to Prof. Jasmin Hefendehl.

## Figures

**Extended Data Figure 1.**
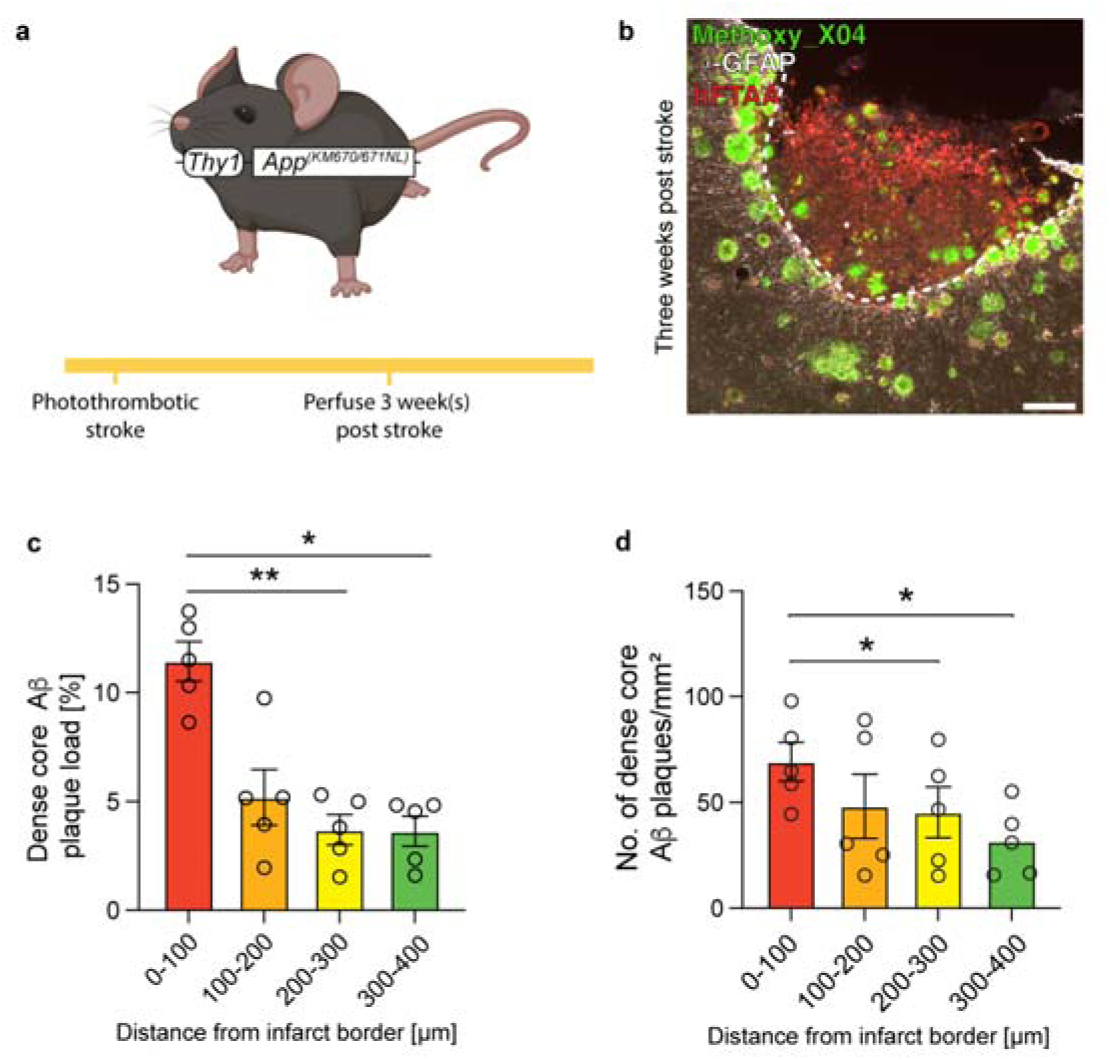
Increased Aβ plaque load in APP23 mice three weeks post stroke. (**a**) Experimental strategy to model ischemic stroke in an alternative mouse model of cerebral amyloidosis (APP23 mice). (**b**) Representative image of an APP23 mouse brain section three weeks post stroke. Scale bar = 200 μm. (**c**) Significantly higher dense-core Aβ plaque load as well as (**d**) the number of dense core Aβ plaques were found in close proximity to the infarct border three weeks post-stroke in APP23 mice (n = 5 mice, 3 males 2 females). Repeated-measures one-way ANOVA with Tukey’s multiple comparison test. * = p < 0.05, ** = p <0.01, *** = p < 0.001, **** = p <0.0001. For full statistical details, see Supplementary Table 2.

**Extended Data Figure 2.**
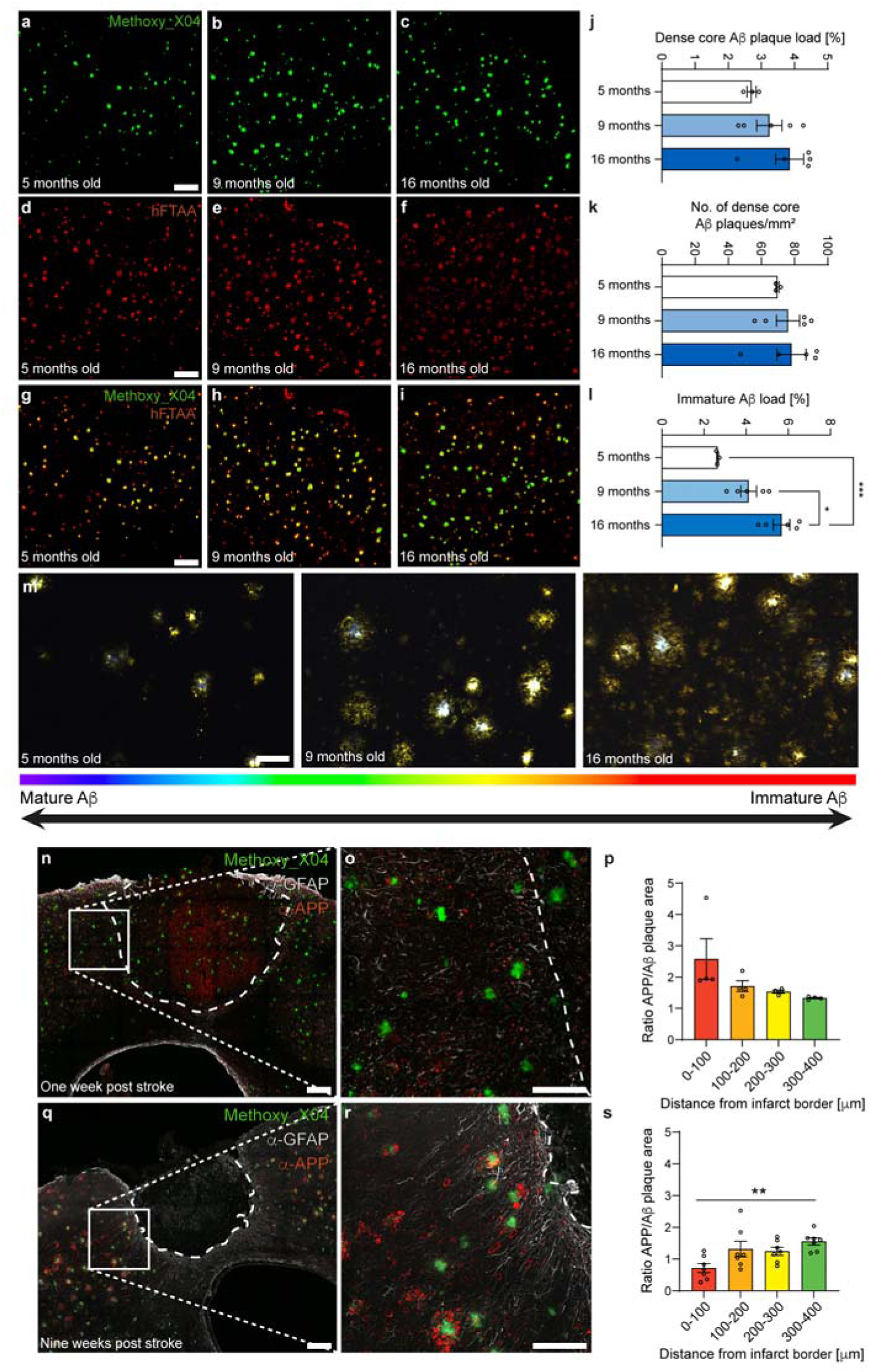
Temporal alterations to Aβ deposits and dystrophic neurite occurrence. (**a-i**) Representative image of an APPPS1 mouse brain section at five, nine and 16 months old. Aβ deposits are labelled with Methoxy X04 (green) and hFTAA (red). (**j**) No significant increase in dense core (Methoxy X04^+^) Aβ plaque load or the (**k**) number of dense core Aβ plaques occurs after five months of age. However, (**l**) there is a significant increase in the hFTAA covered area (immature Aβ load), indicating an increase in immature Aβ species in the brain parenchyma (n = 3-5 mice: 5 months old 2 males 1 female, 9 months old 3 males 2 females, 16 months old 3 males 2 females). (**m**) False-color coding of spectral scan images of APPPS1 brain sections stained with hFTAA and qFTAA taken at five, nine and 16 months old. Note the accumulation of immature (red shifted) Aβ deposits in the brain parenchyma. (**n-p**). No significant difference in Aβ-plaque associated axonal dystrophy was present in close proximity to the infarct border (n = 4 mice, 2 males 2 females) at one week post stroke, however (**q-s**) significantly less Aβ plaque associated axonal dystrophy was present in close proximity to the infarct border (n = 7 mice, 3 males 4 females) at nine weeks post stroke. Dense-core Aβ plaques (n,o,q,r) are visible in green (labelled with Methoxy_X04), glial scar is visible in white (GFAP immunoreactivity), dystrophic neurites are visible in red (amyloid precursor protein (APP) immunoreactivity). (j,k,l) Ordinary one-way ANOVA with Tukey’s multiple comparison test. (p,s) Repeated-measures one-way ANOVA with Tukey’s multiple comparison test. * = p < 0.05, ** = p <0.01, *** = p < 0.001, **** = p <0.0001. For full statistical details, see Supplementary Table 2.

**Extended Data Figure 3.**
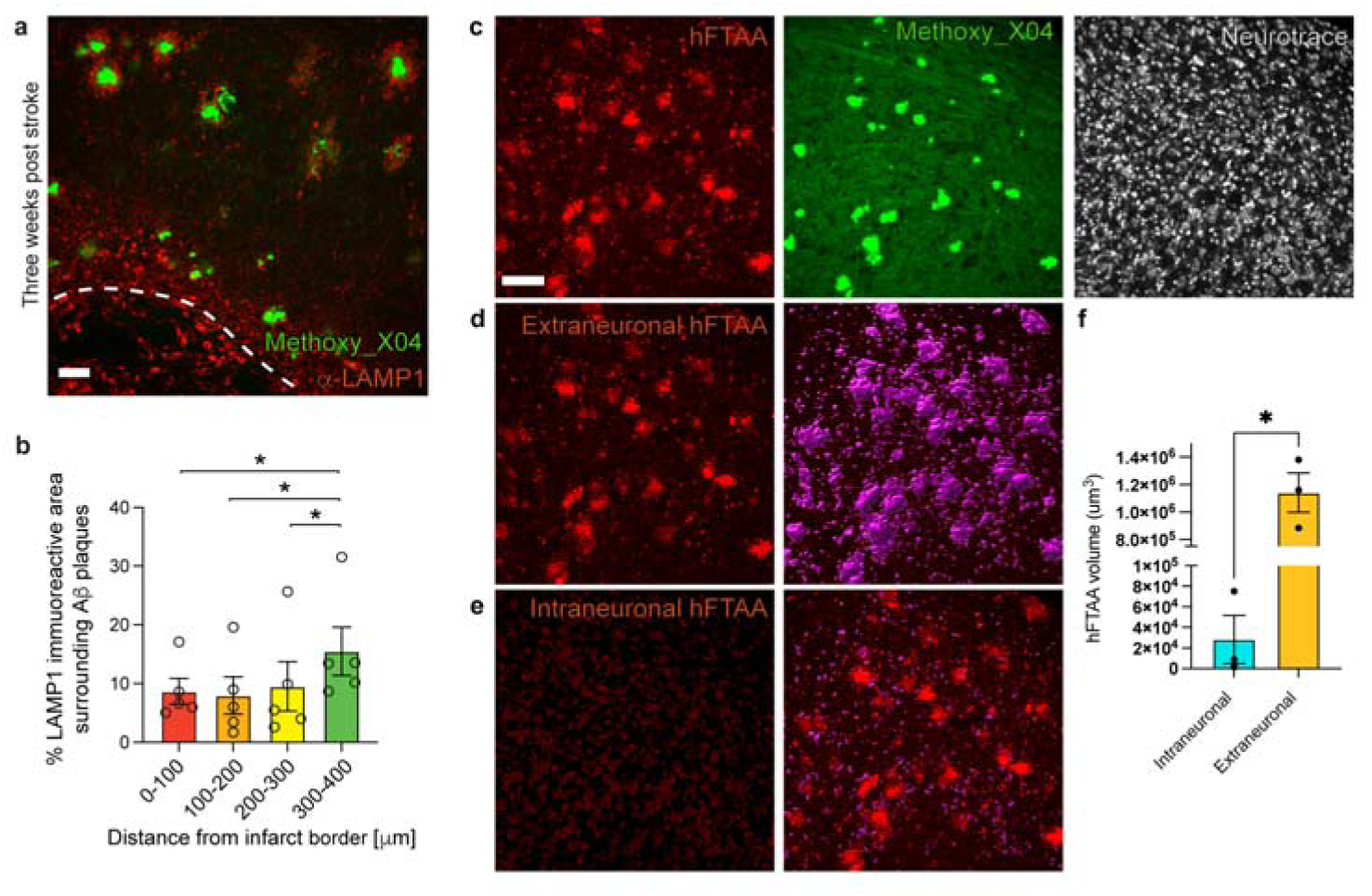
LAMP1 immunoreactivity is decreased proximal to the infarct core. (a) Representative image of LAMP1 immunoreactivity surrounding peri-infarct Aβ plaques. Scale bar = 50 μm. (b) LAMP1 immunoreactivity surrounding Aβ plaques was significantly higher distal to the infarct core (n = 5 mice, 2 males 3 females). (c) Representative images of contralateral hFTAA, Methoxy_X04 and neurotrace at three weeks post stroke. (d) Isolated extraneuronal and (e) intraneuronal hFTAA and corresponding surface reconstruction. Scale bar = 80 μm. (f) Quantification of intraneuronal and extraneuronal hFTAA (n = 3 mice, 1 males 2 females). Repeated-measures one-way ANOVA with Tukey’s multiple comparison test. * = p < 0.05, ** = p <0.01, *** = p < 0.001, **** = p <0.0001. For full statistical details, see Supplementary Table 2.

**Extended Data Figure 4.**
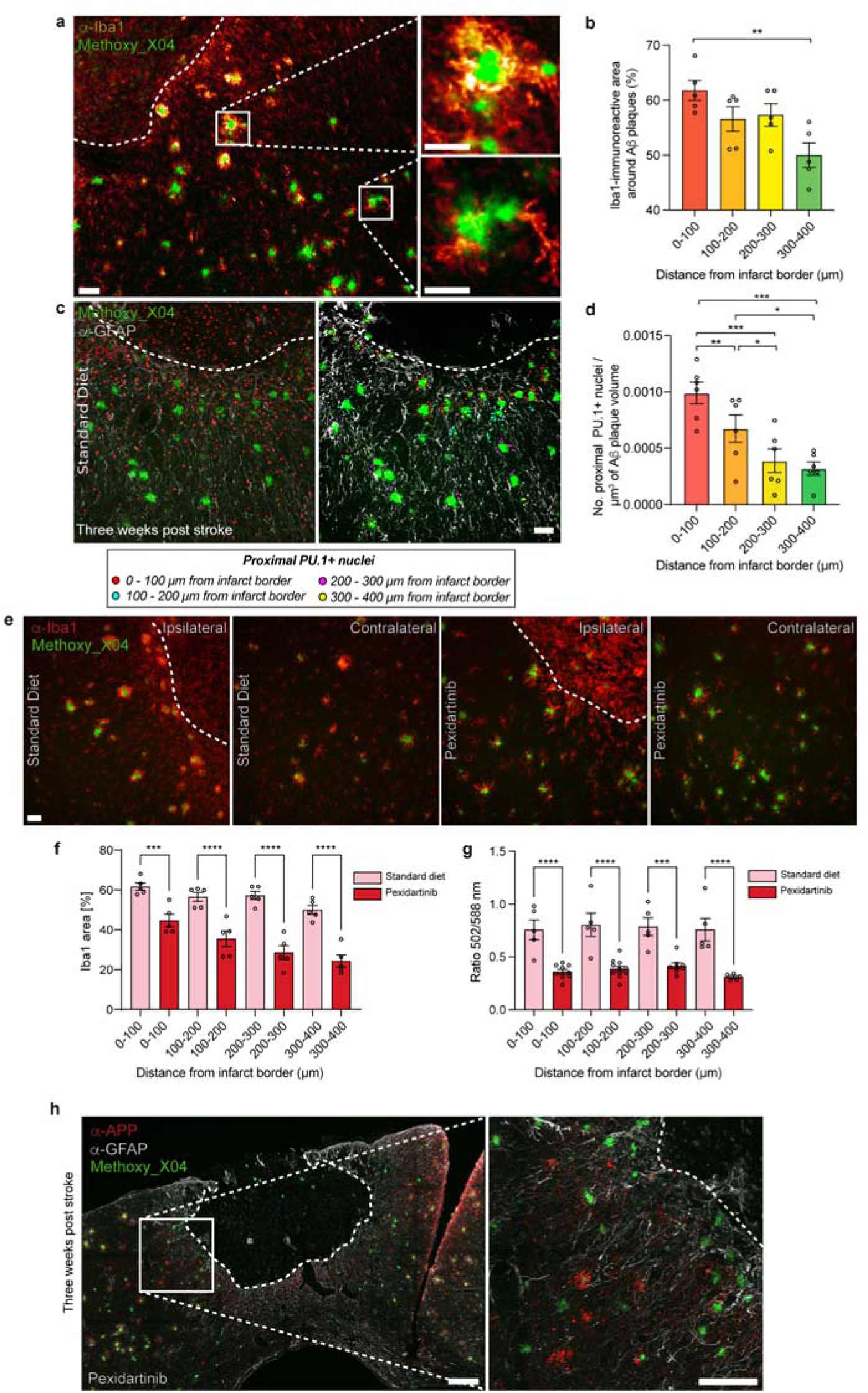
Alterations upon microglia depletion with pexidartinib. (**a**) Representative image of an APPPS1 mouse brain section at three weeks post stroke. Aβ deposits are labelled with Methoxy X04 (green) and microglia are labelled with anti-Iba1 (orange). Scale bar = 50 μm, inset 20 μm. (**b**) Iba1-immunoreactive area around Aβ plaques is significantly higher proximal to the infarct border (n = 5 mice, 3 males 2 females). (**c**, left) Representative image of an APPPS1 mouse brain section at three weeks post stroke. Aβ deposits are labelled with Methoxy X04 (green), glial scar is visible in white (anti-GFAP, white) and microglial nuclei are labelled with anti-PU.1 (red). Scale bar = 20 μm. (**c**, right) Representative image of segmentation procedure used to quantify microglial nuclei around peri-infarct Aβ plaques. (**d**) The number of proximal PU.1^+^ nuclei per μm^3^ of Aβ plaque volume is significantly higher proximal to the infarct border (n = 6 mice, 4 males 2 females). (**e**) Representative images of anti-Iba1 (red) and Methoxy_X04 (green) labelling in the peri-infarct region and contralateral hemisphere three weeks post stroke with or without pexidartinib treatment. Note the reduction of non-Aβ plaque-associated Iba1 immunoreactivity in pexidartinib treated mice compared to standard diet. (**f**) Pexidartinib treatment resulted in a significant reduction in Iba1^+^ area in the peri-infarct region (n = 5 standard diet mice, 3 males 2 females, n = 5 pexidartinib-treated mice, 3 males 2 females) and (**g**) a significant reduction in the qFTAA/hFTAA spectral ratio in the peri-infarct region compared to standard diet three weeks post-stroke (n = 5 standard diet mice, 3 males 2 females, n = 11 pexidartinib-treated mice, 7 males 4 females). (**h**) Representative image of an APPPS1 mouse brain section at three weeks post stroke after six weeks of pexidartinib treatment. Aβ deposits are labelled with Methoxy X04 (green), the glial scar is visible in white (anti-GFAP) and dystrophic neurites (anti-APP) are shown in red. Scale bar = 200 μm. (b,d) Repeated-measures one-way ANOVA with Tukey’s multiple comparison test. (f,g) Ordinary one-way ANOVA with Šidák’s multiple comparison test (f,g).* = p < 0.05, ** = p <0.01, *** = p < 0.001, **** = p <0.0001. For full statistical details, see Supplementary Table 2.

**Extended Data Figure 5.**
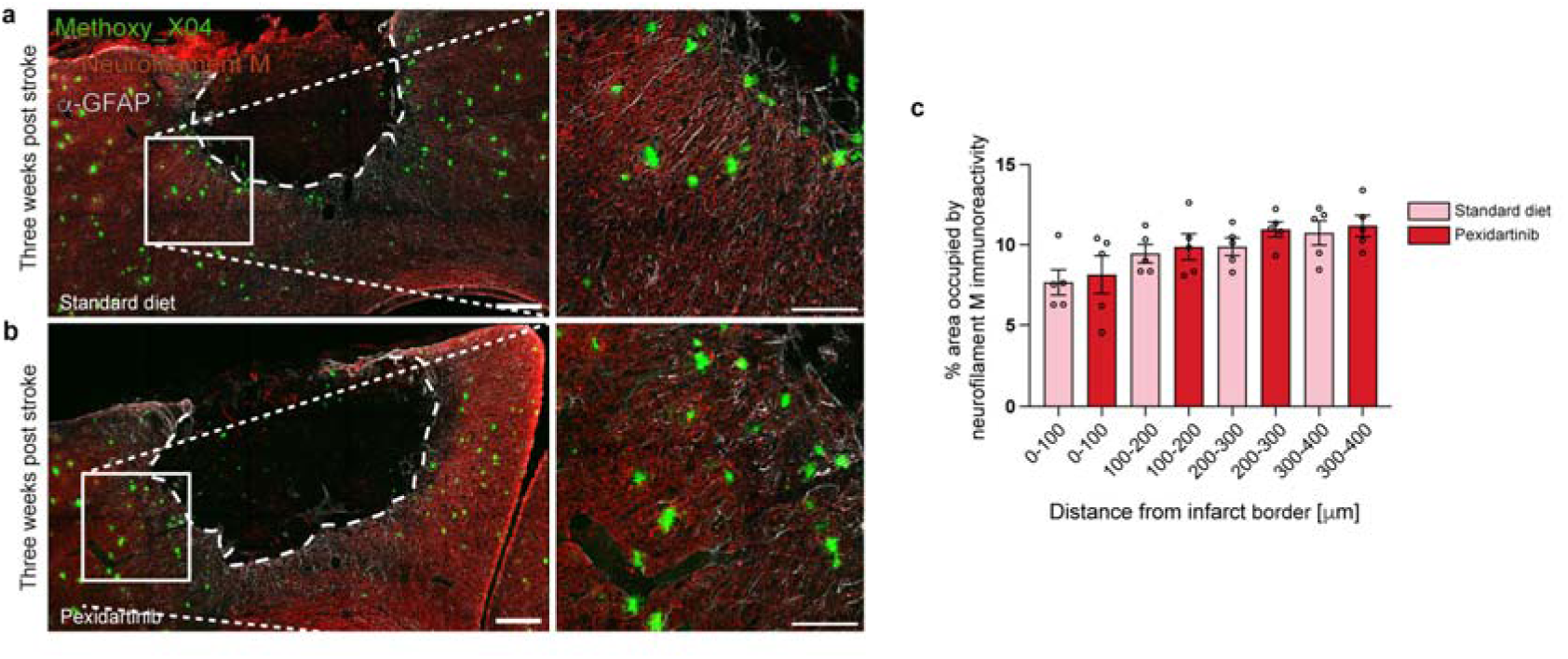
Neurofilament M-occupied area is not significantly different in the presence of pexidartinib. (a) Representative images of neurofilament M immunoreactivity in the peri-infarct region three weeks post stroke from mice either on standard diet or (b) pexidartinib (n = 5 standard diet mice, 3 males 2 females, n = 5 pexidartinib-treated mice, 3 males 2 females). (c) No significant difference was found in the area covered by neurofilament M immunoreactivity between mice treated with pexidartinib or standard diet. Scale bar = 200 μm and 100 μm for insets. Ordinary one-way ANOVA with Holm-Šidák’s multiple comparison test (f,g).* = p < 0.05, ** = p <0.01, *** = p < 0.001, **** = p <0.0001. For full statistical details, see Supplementary Table 2.

**Extended Data Figure 6.**
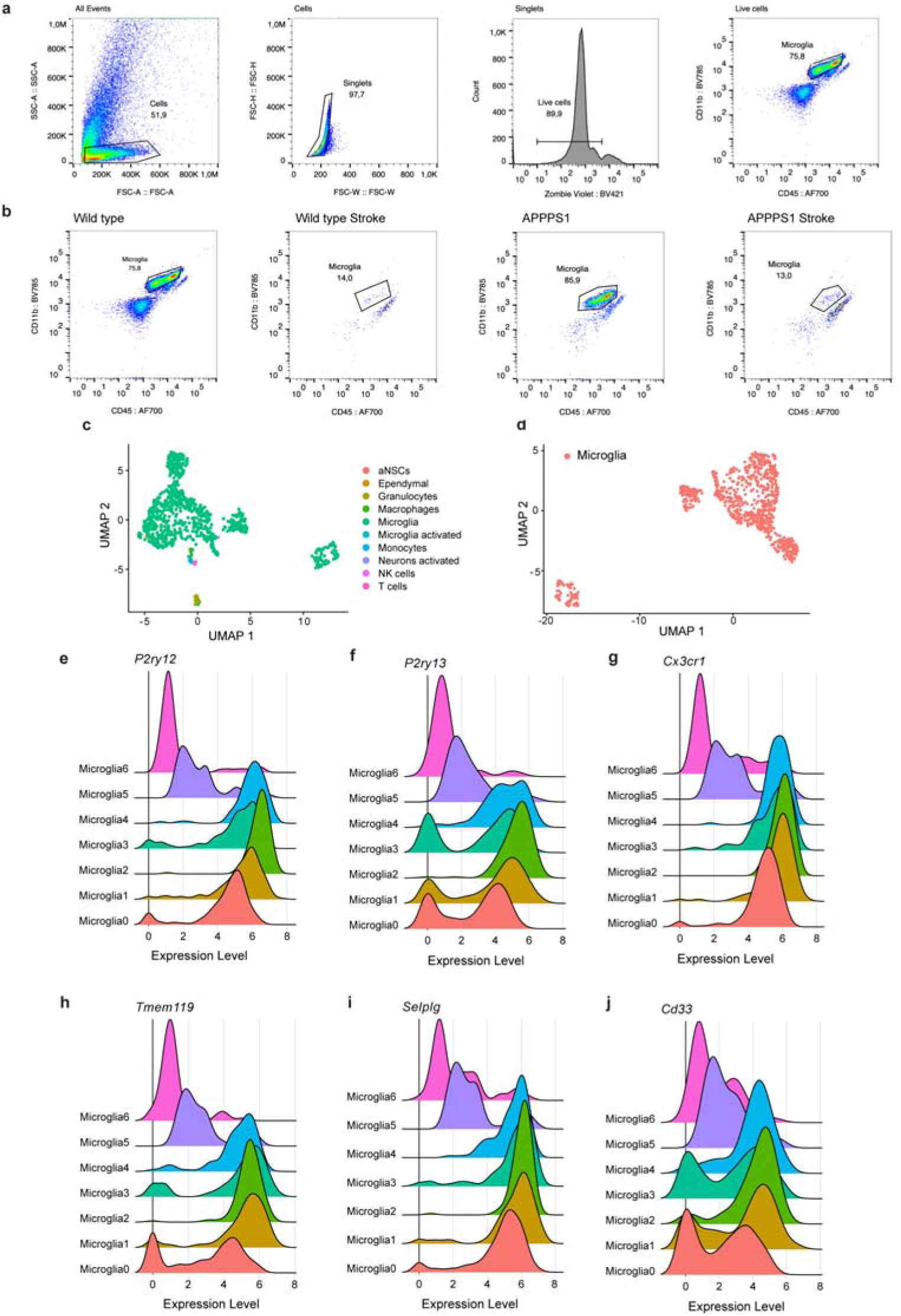
Validation of microglial sorting strategy. (**a**) Representative snapshots illustrating the gating strategy utilized to FACS-sort microglia from (**b**) wildtype, wildtype stroke, APPPS1 and APPPS1 stroke mice. (**c**) UMAP illustrating the results of automatic cell type detection of FACS-sorted microglia prior and (**d**) after exclusion of non-microglial cells. (**e-j**) Ridge plots illustrating the expression level of microglia marker genes within each microglial cluster.

**Extended Data Figure 7.**
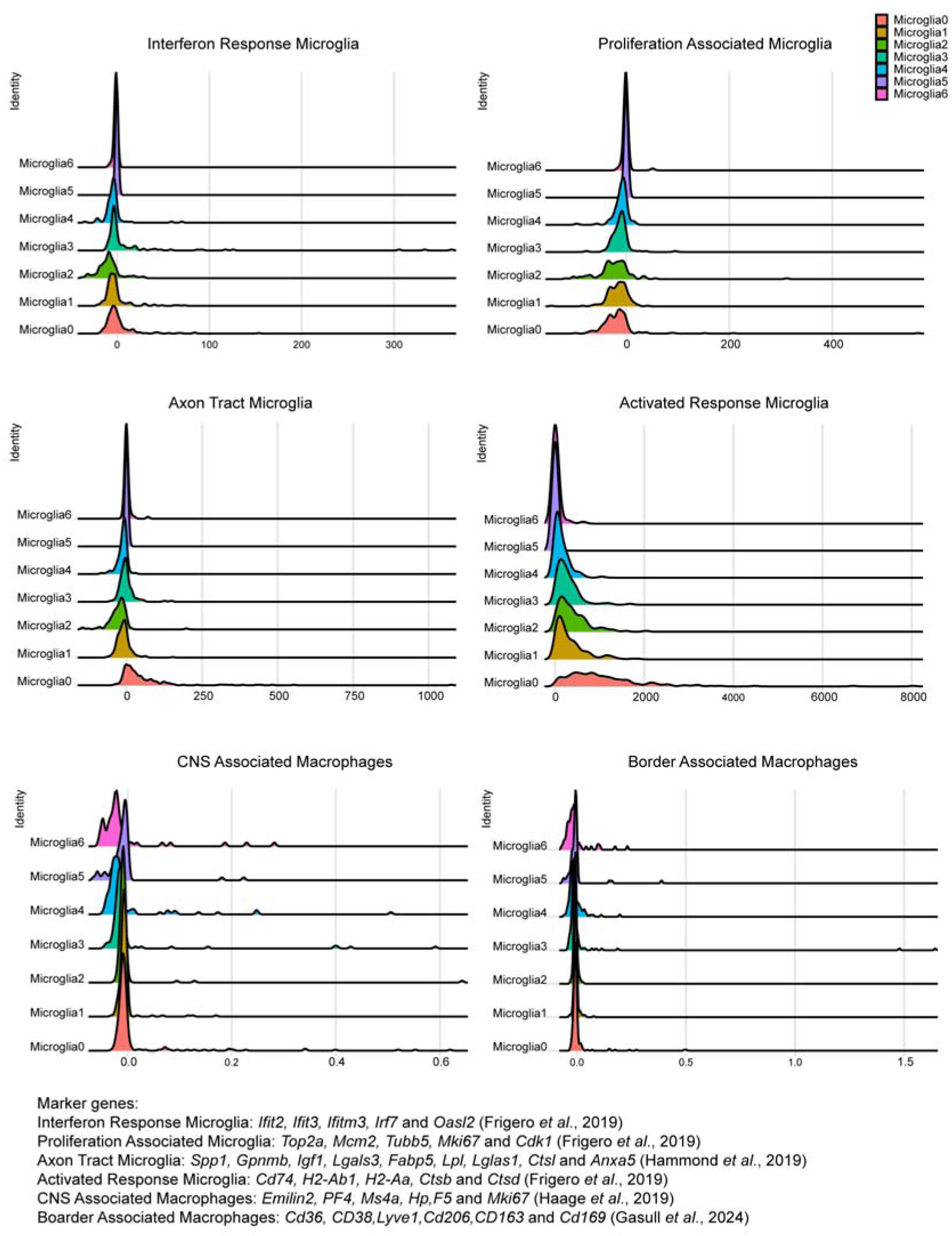
Gene expression profiles of known microglial subtypes. Ridge plots illustrating the expression of marker genes of previously published microglial subtypes.

**Extended Data Figure 8.**
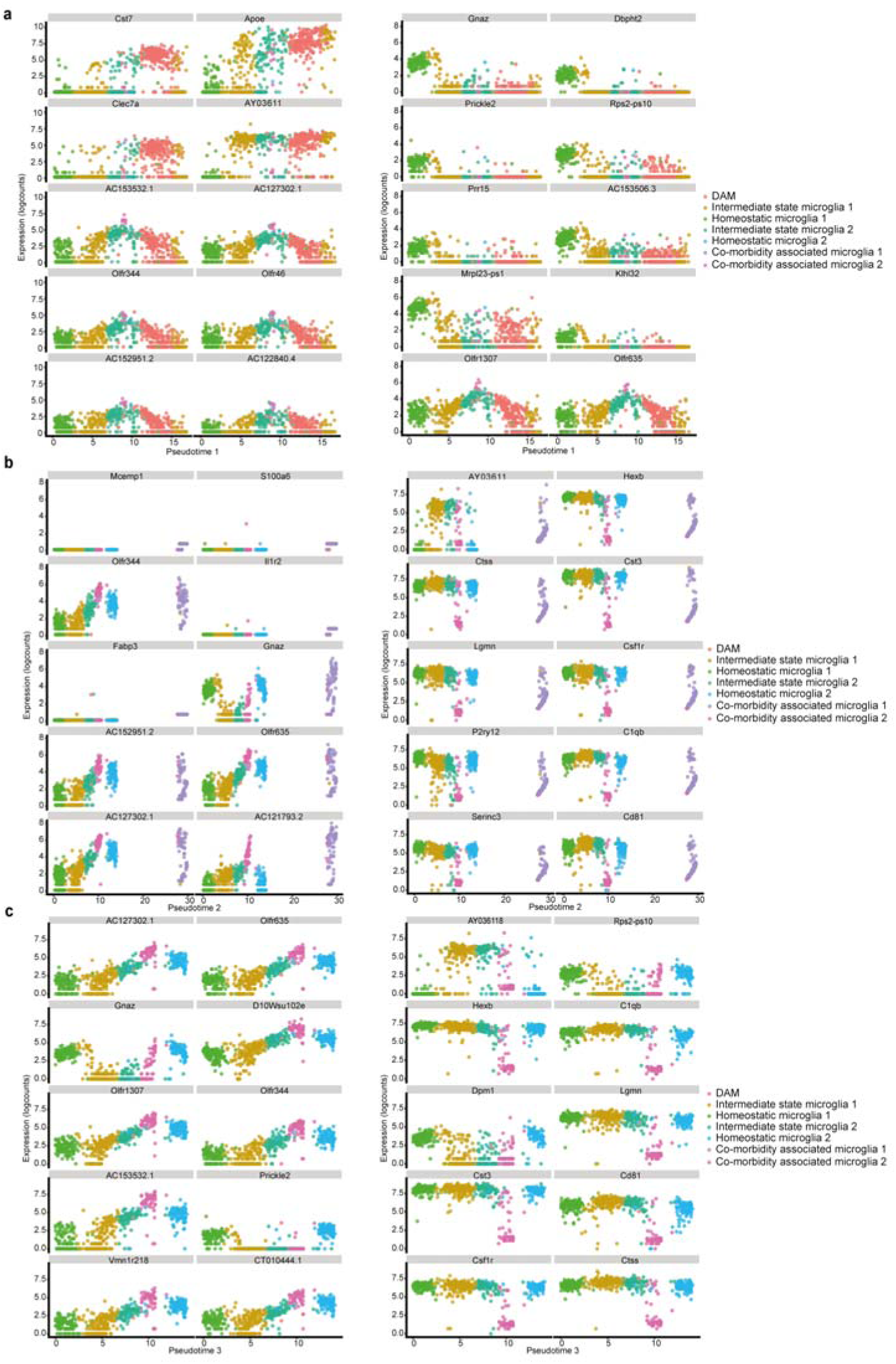
Marker genes determining different pseudotime trajectories. (**a**) Marker genes (upregulated left column, downregulated right column) for the pseudotime trajectories from homeostatic microglia 1 to DAM, (**b**) homeostatic microglia 1 to CoAM and (**c**) homeostatic microglia 1 to homeostatic microglia 2.

## Methods

### Mice (experimental animals)

Animal experiments and husbandry were carried out in accordance with guidelines established by the animal welfare committee of the Johann Wolfgang Goethe-Universität Frankfurt am Main and in accordance with regulations established by the state of Hessen (Germany). Mice were kept under a standard light/dark cycle with access to food and water *ad libitum*. WT and hemizygous APPPS1^1^ transgenic mice harboring two transgenes under the control of the *Thy1* promoter (human APP^KM670/671NL^ and PS1^L166P^) were used in the study. Additionally, hemizygous APP23^2^ transgenic mice (harboring only the human APP^KM670/671NL^) mice were also used. Mice were kept on a C57BL/6J background and were between 5 and 18 months of age. Males and females were mixed for this study. Please see figure legends for n of each sex used in each experiment.

### Ischemic stroke

Mice received an intraperitoneal (i.p,) injection of rose Bengal (Sigma) (100 mg/kg) and were anesthetized using 5% isoflurane. After the surgical anesthetic plane was reached, anesthesia was maintained at 2% isoflurane. A sagittal incision was made in the skin to expose the skull. The targeted brain region (somatosensory cortex, hindlimb) was irradiated through the skull for 20 minutes with 530 nm laser light (11 mW/cm^2^) resulting in thrombosis. Mice were administered a sub-cutaneous injection of carprofen (5 mg/kg) and the incision closed using 4/0 absorbable suture and Histoacryl. Mice were subsequently allowed to recover prior to returning to their home cages.

### Pexidartinib treatment

To deplete brain resident microglia^3^, mice were treated with the drug pexidartinib (PLX-3397, HY-16749 Hyultec GmbH) that was administered via food at a concentration of 290 ppm (290 mg/kg) according to AIN-76A standard diet food of Research Diets Inc. (provided by Ssniff Spezialdiäten GmbH) for three consecutive weeks before the stroke-inducing surgery followed by three more weeks after the stroke.

### Transcardial perfusion and tissue freezing for histology

Mice were sacrificed with an overdose of isoflurane and, shortly after death, transcardially perfused with 20 ml of room temperature (RT) phosphate-buffed saline (PBS) followed by 20 ml of ice-cold 4% paraformaldehyde (PFA). Brains were removed, post-fixed for two hours in 4% PFA at 4 °C and then transferred to 30% sucrose in PBS at 4 °C until sunk. Subsequently, brains were frozen in liquid nitrogen and stored at -20 °C until sectioning. 40 μm free-floating sections were prepared using a sliding microtome (Slee) and subsequently stored in freezing solution (30% glycerol, and 30% ethylene glycol in 1x PBS) at -20 °C until further use.

### Immunohistochemistry/histology

Sections were washed 3x for 5 minutes with PBS and subsequently permeabilized and blocked with 0.5% triton X-100 in PBS and 5% donkey serum (DS) in PBS with 0.02% sodium azide, respectively, for one hour at RT. Sections were then incubated in primary antibodies (Supplementary Table 1) in PBS containing 0.5% λ-carrageenan (Sigma) and 0.02% sodium azide overnight at 4 °C. Subsequently, sections were washed 3x for 5 minutes with PBS with 0.05% tween-20 (PBST) and sequentially incubated with the appropriate secondary antibodies (Supplementary Table 1) in PBS containing 0.5% λ-carrageenan (Sigma) and 0.02% sodium azide at RT for two hours. Sections were then washed 3x for 5 minutes with PBST and transferred to PBS for mounting on Superfrost Plus microscopy slides and coverslipped with Fluoromount-G. For labelling of Aβ using methoxy_X04, a half hour incubation of 2% Methoxy_X04 (Tocris) in DMSO in PBS with 7.66% Kolliphore-EL (Sigma) was performed prior to final washes and mounting.

### Luminescent-conjugated oligothiophene labelling of Aβ *ex vivo*

hFTAA and qFTAA were kindly supplied by Prof. K. Peter R. Nilsson. Sections were washed 3x for 5 minutes with PBS and subsequently permeabilized and blocked with 0.5% triton X-100 in PBS and 5% DS in PBS with 0.02% sodium azide, respectively, for one hour at RT. Sections were then incubated in rabbit anti-GFAP (1:1000, DAKO) in PBS containing 0.5% λ-carrageenan (Sigma) and 0.02% sodium azide overnight at 4 °C. Sections were then washed 3x for 5 minutes with PBST and sequentially incubated with anti-rabbit IgG Alexa 647 (1:1000, Invitrogen) in PBS containing 0.5% λ-carrageenan (Sigma) and 0.02% sodium azide at RT for two hours. Sections were then washed 3x for 5 minutes with PBST and then incubated with (3 μM) hFTAA for 30 minutes, washed 3x with PBST and then incubated with (6 μM) qFTAA for 30 minutes at RT. Sections were then washed 3x for 5 minutes with PBST and transferred to PBS for mounting on Superfrost Plus microscopy slides and coverslipped with Fluoromount-G.

### Luminescent-conjugated oligothiophene labelling of Aβ *in vivo*

Mice were administered an intraperitoneal injection (2 μl/g) of hFTAA in sterile saline (10 mg/ml) one day prior to transcardial perfusion for iDISCO experiments.

### iDISCO

Whole-brain clearing was performed using only the “tissue-clearing” section of the previously described iDISCO protocol^4^ since Aβ labelling was already performed via *in vivo* injection of hFTAA. Briefly, samples were dehydrated in a gradient of methanol in PBS (20%, 40%, 60%, 80%, 100%, 100%) each for one hour on a tube rotator at RT. Delipidation was then performed by incubating the samples in 66% dichloromethane (DCM) with 33% methanol overnight at RT. The next day, samples were transferred to 100% DCM for 45 minutes, following by a 2-hour incubation in dibenzyl ether for clearing. Samples were then transferred into a brown glass vial filled with ethyl cinnamate until imaging. Prior to imaging, brains were transferred to custom 3D printed microscope slides featuring an approximately mouse-brain sized chamber filled with ethyl cinnamate. The chamber was then covered with a glass coverslip and permanently attached to the 3D printed microscope slide with epoxy.

### Wide-field and laser-scanning confocal microscopy

Brain sections were imaged at an epifluorescence microscope (Imager M.1, Zeiss) using a 10x EC Plan-NEOFLUAR 10x/0.3 objective or a laser-scanning confocal microscope (SP8, Leica) using an HC FLUOTAR L 25x/0.95 NA water-immersion objective.

### Spectral scans

For spectral scan analysis of Aβ around stroke lesions, images were taken at a Zeiss 2-photon microscope (Zeiss LSM 880) containing 32 photon multiplier tubes (PMTs) arranged specifically to detect different emission spectra of used luminescent conjugated oligothiophenes (LCOs), such as qTFAA and hFTAA, in a range of 410 – 695 nm in ∼9 nm steps after excitation with a 2-photon laser (Mai Tai® DeepSee™) at 780 nm.

### 2-photon microscopy of cleared mouse brains

iDISCO-cleared mouse brains were imaged using a custom 2-photon microscope (based on a previously published design^5^) equipped with a Chameleon Ultra II laser (Coherent) and controlled by ScanImage software (MBF Biosciences). hFTAA was excited at 900 nm (laser power <50 mW). Emitted light was detected using a non-descanned Hamamatsu detector, a 560 nm dichroic mirror and a 605/70 nm bandpass filter. Z stacks (7 μm steps) were taken using a 10x EC Plan-NEOFLUAR 10x/0.3 objective

### Confocal Raman microscopy

Raman imaging was performed in regions of interest with a confocal Raman microscope Witec 300R^+^ (Witec GmbH, Ulm, Germany) equipped with a 532 nm laser and a 50x objective (NA 0.8) and set to a power of 5 mW in front of the objective. Each scan was acquired in an area of 70x70 µm^2^ with a resolution of 0.333 µm and an integration time of 0.3 s. Stroked APPPS1 mice (1 scan 0-100 μm from the infarct border and 1 scan 300-400 μm from the infarct border) and APPPS1 control mice (1 scan each) were imaged and compared.

### Hyperspectral data processing

Spectral background was subtracted using the ‘shape’ function (filter size: 100) and cosmic rays were removed in the Project FOUR software (Witec GmbH, Ulm, Germany). The data set was then imported to Matlab (The Matlab Inc., Natick, USA) for smoothing using a Savitzky-Golay-filter (window size: 9, order: 3) and normalization using the standard-normal-variate (SNV) method. Spectral unmixing was applied using vertex component analysis (VCA)^6^ the Raman Light App^7^ to identify spectral signatures of Aβ plaques in the tissue. For peak ratio analysis, the β-sheet peak (1238 cm-1) and phenylalanine peak (1005 cm-1) were divided by a peak with a constant intensity (1452 cm-1) in the Aβ spectral signature^8^.

### Data analysis

All statistical analyses and the generation of graphs were performed using GraphPad Prism 9 software. Data analysis was performed using FIJI software for Aβ plaque, APP, Iba1, CD68, and spectral scan analysis. Microglia-Aβ plaque contact-area analysis was performed in tandem with FIJI and Imaris software (Imaris software (v 9.7), Oxford Instruments). Figures were prepared using Fiji, Bitplane Imaris and Adobe Illustrator.

### Spectral scan analysis

For spectral scan analysis, a script was developed by Georgi Tushev from the Max-Planck Institute for Brain Research in Frankfurt am Main, Germany. For spectral scan analysis, Aβ plaques from different conditions were investigated. At least n = 5 Aβ plaques in different regions around stroke (0-400 μm) with varying time points after stroke were examined (1wps, 3wps, and 9wps), as well as the Aβ plaques of microglia-depleted brains that received a stroke (3wps PLX) and Aβ plaques from APPPS1 hemizygous control brains from different age groups (5, 9 and 16 months old).

qFTAA detects mature Aβ fibrils (dense core Aβ plaques), whereas hFTAA detects both mature Aβ and protofibrils^9,10^. ROIs of the Aβ signal were created depending on the emission peaks of qFTAA (502 nm) and hFTAA (588 nm)^11^ using an automated threshold of FIJI software (“Li dark”) and a minimum area of 25 μm² for each Aβ plaque. The ROIs created by the script for the dense-core Aβ plaques and the halos surrounding the Aβ plaques were then assigned to the different regions (0-400 μm) around the lesion. These ROIs were used to assess the Aβ load of dense-core Aβ plaques and diffuse, immature protofibrils by dividing the area of dense-core Aβ plaques and halo in the respective region around the stroke lesion (ratio of dense-core to halo). Furthermore, the maturation state of the Aβ plaques, depending on the signal intensity in each channel captured by the different PMTs, can be assessed. The signal intensity was captured in 32 channels from 410 nm to 695 nm, each covering a signal of around 9 nm of wavelength. The 32 spectra of every Aβ plaque were normalized to the maximum intensity value of this respective Aβ plaque. Afterward, the values for each channel of all Aβ plaques were averaged for each mouse and each region around the lesion.

Two ways of data presentation were chosen to compare the spectra of the conditions and time points in the regions around the lesion. One was to display the averages of all channels as a whole. For the other, the ratios of the peaks of qFTAA (502 nm) and hFTAA (588 nm) were calculated by dividing the value for the qFTAA peak from the value of the hFTAA peak. Statistics were performed using the ratios of the wavelengths.

Furthermore, images absent of stroke lesions (e.g., pictures from APPPS1 control brain sections) were analyzed as described above without any distance measurements. Notably, GFAP was used to identify the lesion border, which was detected with a secondary antibody conjugated to Alexa-647. Faint signals of this secondary antibody may be visible in the pictures and hence in the spectra of 647 nm. This signal, however, does not influence the ratio peak analyses of qFTAA and hFTAA.

### Image stitching

Several pictures covering the lesion and peri-infarct region taken at the epifluorescence microscope required stitching to ensure a covered area of 400 μm tissue around the lesion in one image. This was accomplished using the 2D stitching plugin provided by FIJI software. When imaging with SP8 confocal microscope to cover sufficient tissue around the lesion, tile scans were performed, and the single images were subsequently stitched automatically to a merged image by the provided Leica LAS X software.

### Regions around lesion

To analyze the peri-infarct region around the lesion for images taken at the epifluorescence microscope, the SP8 confocal microscope, or the Zeiss 880 2-photon microscope, the peri-infarct region was segmented into regions of 100 μm width starting from the delineation of the glial scar up to 400 μm distal from the lesion. Using FIJI software, along the border of the lesion, a ROI was manually drawn with the glial scar as a template. This ROI was enlarged by the respective size to get regions containing all the tissue for up to 400 μm. (lesion-100 μm, lesion-200 μm, lesion-300 μm and lesion-400 μm). The ROIs were then subtracted from each other appropriately to atain the desired region ROIs (0-100 μm, 100-200 μm, 200-300 μm, and 300-400 μm).

### Aβ plaque analysis

Epifluorescence images were processed with FIJI software to analyze the Aβ plaques surrounding the lesion in the different regions or on control sections. The background of the Methoxy-X04 signal pictures was subtracted with the rolling ball method with 50 pixels, followed by blurring according to the Gaussian blur filter (sigma = 2). A threshold was applied according to the mean grey value of the image plus one standard deviation that was measured using FIJI excluding the infarct lesion. ROIs for the plaques were created using the “Analyze particles” function. Aβ plaques were selected manually depending on the distance from the lesion. With the information provided by the FIJI software, the Aβ plaque number/mm², Aβ plaque area/mm² (plaque load), and the average Aβ plaque size were calculated. For Aβ plaque analysis of control sections, all Aβ plaques visible in the field of view were considered (plaques touching the edges were excluded using the “analyze particles” function). Aβ plaque analysis was performed as described above to determine diffuse Aβ in control sections using Methoxy-X04 and hFTAA. The determined load for both dyes was subtracted from one another to attain the diffuse Aβ load.

### APP analysis

8-bit maximum projection pictures of the acquired merged SP8 confocal images were used for analysis. Aβ plaque analysis was performed similarly as described above using the pictures of the channel containing the Aβ plaque signal. A threshold with the mean grey value plus 2*standard deviation (values obtained excluding the lesion) was applied for the Aβ plaques. As only the total Aβ plaque area per region was needed, any signal of plaques ranging outside of one of the regions was cut out and removed to only receive the total Aβ plaque area per region. The Aβ plaque area per region around the lesion was automatically calculated by FIJI software.

The fluorescent background of images with APP signal was subtracted using the rolling ball method provided by FIJI software with a size of 50 pixels. Afterward, for analysis of the APP area / Aβ plaque area ratio, a threshold was applied with mean grey value plus 2*standard deviation of the whole image, excluding the infarct lesion. The area of the APP signal was measured in all regions using the “Analyze Particles” function, and the total APP area per total Aβ plaque area for the different regions was calculated.

### Iba1 analysis

Aβ plaque load analysis was performed as described above on 8-bit maximum projection images acquired via confocal microscopy. The ROIs for the plaques were enlarged to obtain ROIs covering 15 μm around the plaques. All ROIs were saved for later analysis. The background of the channel containing the Iba1 signal was subtracted with the rolling ball method with a size of 50 pixels. All the Iba1 signal outside the region of interest and 15 μm around the Aβ plaques was removed only to analyze the microglia close to the Aβ plaques and in the respective region around the lesion. An automated threshold provided by FIJI (“Huang2”) was applied, and by using the “Analyze Particles” function, the covered area of the Iba1 signal around the Aβ plaques was determined.

### Microglia-Aβ plaque contact area analysis

Before analyzing the microglia-Aβ plaque contact area with Imaris software, using FIJI software, the 16-bit z-stacks acquired from confocal imaging, additional channels containing the different regions around the lesion (0-400 μm) in different colors were created. These channels were later used to assign the Aβ plaques to the regions in Imaris.

Then, using Imaris software, surfaces were reconstructed using the Methoxy-X04-positive signal to create surfaces for Aβ plaques and the Iba1-positive signal to create surfaces for microglia. Aβ plaque surfaces were manually selected and deleted to leave only Aβ plaques from one region around the lesion per reconstructed Aβ plaque surface. Microglial surfaces were generated based on the Iba1 signal in the entire image. The area of Aβ plaque surfaces in contact with microglial surfaces was calculated using the “XTension” “Surface-Surface Contact Area”. For this, the Aβ plaque surfaces were selected as the primary surface and the microglial surface as the secondary surface.

### CD68 analysis

For analyzing CD68 intensity in microglia close to Aβ plaques, Aβ plaque load analysis and Iba1 analysis were performed as described above. The ROIs for Iba1-positive microglia were saved for each region around the lesion. Using the 8-bit max projection pictures of the channel showing the CD68 signal and the ROIs acquired for the Iba1 positive signal, CD68 intensity (mean grey value) was measured in microglia near Aβ plaques using FIJI software.

### PU.1 analysis

For analyzing PU.1 + nuclei in close proximity to Aβ plaques, a surface was created in Imaris for each channel (Methoxy_X04 and PU.1) in the peri-infarct regions of interest (i.e. 0-100 µm, 100-200 µm, 200-300 µm and 300-400 µm). A “distance to image border” filter of minimum 10 was set for the Methoxy_X04 surface to exclude Aβ plaques close to the image border. For the PU.1 surface, “spots” were created with an estimated XY diameter of 5 µm with background subtraction. Spots that were more than 50% outside of the image border of each ROI were excluded. The Xtension “find spots close to surface” with a threshold of 10 µm was subsequently used to detect PU.1 nuclei in close proximity to Aβ plaques.

### Neurofilament-M analysis

The background of 8-bit maximum projections of confocal images containing the neurofilament-M signal was subtracted with the rolling ball method with 10 pixels, and subsequently, a threshold was applied according to the mean grey value of the whole image plus one standard deviation that was measured using FIJI not considering the infarct lesion. The total area of the present neurofilament-M signal was measured for each region up to 400 μm around the lesion using the “Analyze Particles” function.

### LAMP1 analysis

8-bit maximum projections of SP8 confocal Z-stack images were used for analysis. Aβ plaque analysis was performed in FIJI as described above. A gaussian blur with a sigma of 2 was then applied to the LAMP1 channel. For analysis of the LAMP1 area, a threshold was applied with mean grey value plus 2*standard deviation of the whole image, excluding the infarct lesion. ROIs of Aβ plaques were expanded by 20 μm to analyze Aβ plaque-associated LAMP1 immunoreactivity The percentage of area occupied by LAMP1 immunoreactivity was measured within the expanded Aβ plaque ROIs for the different regions using the “measure” function.

### P2RY1 analysis

8-bit maximum projection pictures of SP8 confocal Z-stack images were taken of three weeks post stroke brain sections labelled with anti-P2RY1, anti-Iba1 and Methoxy_X04. In Imaris, surfaces were generated of the microglia and Methoxy_X04. Additionally, a surface was manually generated for the infarct core. Firstly, using the filter function in Imaris surfaces, the microglia surfaces were separated into ROIs radiating from the infarct core (1-100 μm, 101-200 μm, 201-300 μm, 301-400 μm). These were again filtered to attain microglia within 20 μm distance from the Methoxy_X04 surfaces within the respective regions. Using these surfaces of peri-infarct Aβ plaque-associated microglia within the respective regions, the mean intensity of P2RY1 was compared. For contralateral images, the same procedure was followed however excluding the stroke regions therefore attaining the mean intensity of P2RY1 within peri-infarct Aβ plaque-associated microglia on the contralateral side.

### Intra/extra neuronal hFTAA signal segmentation

8-bit maximum projection pictures of SP8 confocal Z-stack images were taken of three weeks post stroke brain sections labelled with Neurotrace 500/525 (ThermoFisher, N21480), hFTAA and Methoxy_X04. The following channels were aquired; Methoxy_X04 in blue, neurotrace, hFTAA and methoxy_X04 in green and hFTAA alone in red. Surfaces were generated in Imaris for Methoxy_X04 and hFTAA. Due to the overlapping emission spectra of neurotrace, hFTAA and Methoxy_X04 in green, the hFTAA surface was used as a mask to set all voxels of the neurotrace/hFTAA/Methoxy_X04 channel within the hFTAA surface to zero; thus generating a new channel with only neurotrace signal (i.e. removing Aβ plaques from this channel). A surface was then generated of the isolated neurotrace signal which was used as a mask on the hFTAA channel to set voxels outside to zero to create an intraneuronal hFTAA channel. This was repeated again, setting the voxels inside to zero to create an extraneuronal hFTAA channel. However, this extraneuronal hFTAA channel still contains Methoxy_X04+ dense core Aβ plaques (which are also hFTAA positive). Therefore it was necessary to remove the Methoxy_X04 labelled Aβ signal from the extraneuronal hFTAA channel to attain a new channel containing Aβ that is exclusively labelled with hFTAA. Therefore, the surface of the Methoxy_X04 channel was used as a mask to set inside voxels of the extracellular hFTAA channel to zero resulting in an extraneuronal hFTAA channel without methoxy_X04+ dense core Aβ plaques. Surfaces were then generated of the extraneuronal hFTAA channel minus methoxy_X04 and the intraneuronal hFTAA channel to attain the total volume of extraneuronal and intraneuronal hFTAA within the image.

### Statistical analysis

Statistical analyses were performed using the software Prism 9 (GraphPad Software). In general, for all the conducted experiments and analyses, when comparing the results for the regions around the lesion (0-400 μm) within one condition (1wps, 3wps, 9wps, or 3wps pexidartinib), they were evaluated using a repeated-measure one-way ANOVA Tukey’s multiple comparisons test unless otherwise stated. There, sphericity was assumed, and the mean of each region was compared to the mean of every other region. A mixed-effect analysis was performed if values were missing for the repeated measures analysis.

For multiple comparisons between the different conditions (for example, comparing 0-100 μm region around the stroke between 1wps, 3wps, 9wps, and ctrl; or comparing the different age groups (5, 9, 16 months) of control sections) an ordinary one-way ANOVA Tukey’s multiple comparisons test was performed. For comparisons of the four regions around the lesion of 3 wps standard diet to 3 wps pexidartinib treatment, an ordinary one-way ANOVA Šídák’s multiple comparisons test was performed with selected comparisons only to compare the respective regions (e.g., 0-100 μm SD vs 0-100 μm PLX). Possible outliers were identified using the “Identify Outliers” function of Prism 9 using the ROUT method. Cleaned data were then further processed as described above. Unless otherwise stated, graphs display the individual values and the mean ± standard error mean. For details regarding statistical tests see Supplementary Table 2.

### scRNA-seq

Mice received a photothrombotic stroke three weeks before microglial isolation. Mice that suffered from a photothrombotic stroke and control APPPS1 and wild-type animals were deeply anesthetized with ketamine/xylazine solution and perfused with ice-cold 1x PBS. The brains were cut out, and the right cortex was dissected. The stroke lesion was dissected, including a small amount of tissue surrounding the lesion. The control animals’ whole cortex of the right hemisphere was dissected.

The dissected tissue was homogenized in a petri dish covered with 500 μl dissection medium (0.5 % D-Glucose and 0.1 mg/ml DNase I in 1x HBSS w/o Ca+Mg) by mincing it into small pieces with a scalpel. The tissue was transferred into a 7 ml capacity Dounce homogenizer along with a 1 ml ice-cold dissection medium. The solution containing tissue (around 2 ml total volume) was slowly homogenized five to six times and transferred into a 5 ml Dounce homogenizer, followed by three additional slow homogenization steps. The solution was transferred through a 70 μm cell strainer into a 15 ml Falcon® tube, and the cell strainer was washed with 1x HBSS and filled up to 15 ml total volume. Afterward, it was centrifuged for 20 minutes at 300 G at 4 °C, and no brake was set in the centrifuge. 4 ml of 30 % isotonic Percoll in HBSS was added into a fresh 15 ml Falcon tube and underlaid with 4 ml 37 % Percoll, additionally dyed with Phenol red to differentiate the different Percoll gradients. The supernatant of the tissue solution was discarded, and the cell pellet was resuspended in 5 ml 70 % Percoll and put underneath the 37 % Percoll using a long glass Pasteur pipette. The solution was centrifuged at 800 G and 4 °C for 30 minutes with no brakes set in the centrifuge. After centrifugation, the myelin layer was first carefully removed from the top of the gradient. The cells forming a white halo from the interphase between 37 % and 70 % gradients were carefully collected and transferred into a new 15 ml Falcon tube that was then filled with FACS buffer (10 mM EDTA, 5 mM HEPES and 2% FCS in 1x HBSS w/o Ca + Mg). The tube was inverted and flicked to dissolve possible remaining high-density Percoll at the bottom of the tube and centrifuged at 300 G and 4 °C for 20 minutes. Afterward, the supernatant was discarded, and the cell pellet was resuspended in 200 μl FACS buffer and transferred into a 1.5 ml Eppendorf® cup. 0.5 μl Fc block (1:500, BD, Art. No. 553141) was added to the solution and incubated for 10 minutes at 4 °C on a rotation wheel. It was washed with 1 ml FACS buffer and centrifuged at 300 G and 4 °C for five minutes with the brake off. The supernatant was discarded and ideally reduced to a total volume of around 100 μl. Then, 0.5 μl of each CD11b – BV785 (1:200, Biolegend, Art.No. 101243) and CD45 – AF700 antibody (1:200, Biolegend, Art.No. 103128)was added to the solution and incubated for 15 minutes at 4 °C on the rotation wheel in the dark. The solution was washed again with 1 ml FACS buffer and centrifuged for five minutes at 300 G and 4 °C with brakes off. The supernatant was discarded and filled up to 250 μl with FACS buffer.

With a cell sorter (Sony SH800S cell sorter), single cells positive for CD11b and CD45 were sorted into 384-well plates containing lysis buffer and oligonucleotides for Smart-seq2^12^ and stored at -80 °C until further processing.

Sorted cells were further processed using an adopted Smart-seq2 protocol for 384-well plates^13^. In short, 384-well plates containing single cells were thawed and cells were lysed and mRNA denatured by incubating the plates at 95°C for 3 minutes and put back on ice. Subsequently, 2.7 μl of 440 first-strand cDNA reagent mix, containing 1x First-strand buffer, 5 mM DTT, 1 M betaine, 14 mM MgCl2, 5 U RNase inhibitor (Takara), 25 U Superscript II Reverse Transcriptase, and 1 μM Template-Switching oligonucleotides in nuclease-free water, was added to each well. Plates were incubated at 42°C for 90 minutes, 70°C for 15 minutes, and kept on hold at 4°C. Afterward, 7.5 μl of the pre-amplification mix, consisting of 1x KAPA HiFi HotStart Readymix and 0.12 μM ISPCR primers in nuclease-free water, was added to each well. Plates were then incubated as followed: 98°C for 3 minutes, 23 cycles of 98°C for 20 seconds; 67°C for 15 seconds; and 72°C for 6 minutes, then at 72°C for 5 minutes and kept on hold at 4°C. Purification of the resulting cDNA was performed using SeraMag SpeedBeads containing 19% w/v PEG 8,000 and a 1:0.8 ratio of cDNA/beads was used for cDNA precipitation. Purified cDNA was eluted in 14 μl of nuclease-free water. The quality of cDNA was randomly checked in 5% of the wells using the Tapestation High-Sensitivity D5000 assay. For the tagmentation reaction, 0.5 μl of 50-150 pg cDNA was mixed with a 1.5 μl Tagmentation mix containing 1x Tagment DNA buffer and Amplicon Tagment mix (Nextera XT DNA sample preparation kit). Plates were incubated at 55°C for 8 minutes and kept on hold at 4°C. 0.5 μl Neutralize Tagment buffer was added and incubated at room temperature for 5 minutes to inactivate Tn5. PCR amplification of adapter-ligated cDNA fragments was performed in a final volume of 5 μl containing the two index primers (Nextera XT Index kit) in a 1:5 dilution and Nextera PCR master mix. Plates were then incubated as followed: 72°C for 3 minutes, 95°C for 30 seconds, then 14-15 cycles of 95°C for 10 seconds; 55°C for 30 seconds; and 72°C for 30 seconds, then 72°C for 5 minutes and kept on hold at 4°C. 384 wells (the entire plate, including negative controls) were pooled in a single 2 ml tube (Eppendorf). Purification of cDNA was performed by adding 400 μl of cDNA in a 1:1 ratio with SeraMag SpeedBeads in 19% w/v PEG 8,000 in a 1.5 ml LoBind tube. Beads were washed with 1 ml of 80% ethanol. Purified cDNA was eluted in 200 μl of nuclease-free water. The concentration of the library was measured by Qubit according to the manufacturer’s instructions. The size distribution of the library was measured using the Tapestation High-Sensitivity D1000 or D5000 assay. The cDNA library pool was stored at -20°C until ready for sequencing.

The cDNA library pool was diluted to 2 nM and prepared for sequencing using the NextSeq 500 High Output Kit v2 or NovaSeq 6000 SP reagent kit 1.5 (75 cycles) according to the manufacturer’s instructions. Sequencing was performed on an Illumina NextSeq 500 or Illumina NovaSeq 6000 instrument. Preprocessing of the data was performed as previously described^13^.

Analysis was performed using the Seurat package and Harmony pipeline with R software. For the detailed cluster analysis we excluded any transcripts coded on either of the sex chromosomes, or do not have any canonical name yet (i.e. cDNAs from the RIKEN project or Gm* Genes). Samples were integrated across batches based on the 2000 most variable genes using the integrated anchoring approach as implemented in Seurat. Reads were Normalized and SCT-Transformed. UMAP dimensions were calculated based on the first 20 dimensions of the Harmony data reduction (selection based on elbow criteria of the skeeplot). Initial clustering was performed with a resolution of 0.6. Cell Cycle scoring was based on tinyatlas mouse gene data (https://github.com/hbc/tinyatlas). For identification of cell types and confirmation of microglial identity was based on a publicly available scRNA data^14^ and implemented in the celdex package. For any further analyses only cells labelled as Microglia were included. After gene and cell filtering, reclustering across all samples was performed including optimization for the clustering resolution (i.e. 0.6): A range from 0.1 to 1.2 resolution was implemented and best clustering was selected based on the most stable condition using clustree plots. A total of 7 clusters (Microglia 0-6 was identified). DAM scores and Homeostatic scores were calculated (AddModuleScore) on gene-sets previously published^15^. Scores were distributed bimodally and the cutoff for DAM+ or Homeo+ cells was manually set at the lowest point of inflection between the two peaks. Differences in cell distribution were tested using chi-square tests, and pairwise chi-square test with post hoc correction. For identifying markers and differentially expressed genes between two conditions the FindMarkers function with the “LR” method was applied, adjusted p-values are reported. GO-term enrichment analysis was done with the profiler package with the universe of all genes detected in the scRNA sample as background reference and the SCS-significance threshold as published^16^. Pseudotime analysis was performed using the slinghot packages with Microglia 2 as the starting cluster and default conditions. TSCANs function testPseudotime identified Genes differentially expressed across trajectories.

### Brain collection for spatial transcriptomics

Mice were sacrificed three weeks post-stroke by cervical dislocation, rapidly decapitated and the brain removed and rolled on a tissue paper (VWR) to remove excess blood. The brain was then transferred to a sterile Petri dish and cut into two hemispheres using a new glass coverslip. Each hemisphere was then transferred to a Peel-a-Way paraffin embedding mold filled with Optimal Cutting Temperature (OCT, Leica Biosystems) compound (from a freshly-opened bottle). The embedding molds were then transferred to a Coplin jar filled with isopentane, which was the placed in a box containing dry ice and ethanol. To determine the RIN of each brain sample, 10x 10 μm coronal sections were taken from the OCT block containing the contralateral hemisphere using a freezing microtome (Leica Biosystems), whereas the OCT blocks containing each ipsilateral side were used exclusively for spatial transcriptomics.

### RNA integrity number (RIN) check

The RNA was extracted using RNeasy Mini Kit (Qiagen, USA) according to the manufacturer’s protocol. The quality of the tissue was then checked by calculating its RNA Integrity Number (RIN), using the Agilent RNA 6000 Nano Assay kit with a 2100 Bioanalyzer (Agilent, USA), following the manufacturer’s protocol, as suggested by the Visium Spatial Tissue Optimization User Guide (CG000240 Rev E, 10x Genomics). Samples only with a RIN ≥ 7 were eligible for the transcriptomics experiments.

### Spatial transcriptomics

Spatial transcriptomics were carried out using the Visium platform (10x Genomics) following the manufacturer’s instructions. Briefly, the slide was processed by following the methanol fixation, Immunofluorescence Staining & Imaging for Visium Spatial Protocols (CG000312, 10x Genomics). Briefly, the Visium Spatial Gene Expression slide was fixed, blocked, and then stained with primary antibodies for 30 minutes at room temperature, washed five times, stained with secondary antibodies for 30 minutes at room temperature, and then washed another five times. The slide was mounted using RNase-free glycerol, and imaged using Zeiss Cell Observer Z.1 microscope. After imaging, the slide was further processed by following the Visium Spatial Gene Expression Reagent Kits - User Guide (CG000239, 10x Genomics).

RNA was extracted according to manufacturer’s instructions (Visium Spatial Gene Expression Slide & Reagent Kit, 16 rxns PN-1000184).The Library preparation was conducted according to the manufacturer indications (Visium Spatial Gene Expression Slide & Reagent Kit, 16 rxns PN-1000184).

The Raw reads were aligned against the mouse genome (mm10) and counted by StarSolo^17^ followed by secondary analysis in Annotated Data Format. Preprocessed counts were further analyzed using Scanpy^18^. Basic data quality control was conducted by taking the number of detected genes and mitochondrial content into consideration as well as coverage by tissue on spots. We removed 10502 data points in total that did not express more than 1000 genes or had a mitochondrial content greater than 30% and were not covered by tissue. Furthermore, we filtered 39771 genes if they were detected in less than 30 cells (<0.01%). Raw counts per cell were normalized to the median count over all cells and transformed into log space to stabilize variance. We initially reduced dimensionality of the dataset using PCA, retaining 50 principal components. Subsequent steps, like low-dimensional UMAP embedding and cell clustering via community detection, were based on the initial PCA.

Final data visualization was done using the CZ CELLxGENE Annotate package in combination with the CellxGene VIP extension^19^.

### Human brain section processing and analysis

Mounted human brain sections were sections were heated at 58°C for 1 h, then deparaffinized and rehydrated (3 x 10 min Xylol, 3 x 5 min 100% EtOH, 2 x 5 min 96% EtOH, 1 x 5 min 70% EtOH, 1 x 5 min ddH2O)using standard protocols. For antigen retrieval, sections were boiled in citrate buffer (1.8 mM citric acid, 8.2 mM trisodium citrate, pH 6) at 90°C for 30 min. Nuclear staining was then performed by incubation with nuclear fast red (Morphisto, Art. No. 10264), according to manufacturer’s instructions. Subsequently, non-specific antibody binding was blocked by incubation with 5% normal donkey serum (in 0.3% Triton X-100 in PBS) for 1 h, followed by primary antibody incubation (goat anti-Iba1, 1:600, Novus, Art. Number NB100-1028) in 2% serum (in 0.3% Triton X-100 in PBS), over two nights at 4°C. After washing, the sections were incubated with the secondary antibody (donkey anti-goat, AF647-conjugated, Jackson Immuno Research, Art. Number 705-605-147, 1:250 in PBS) for 2 h at RT. For the LCO staining, sections were incubated with qFTAA (2.4 µM in PBS) and hFTAA (0.77 µM in PBS), according to standard protocols^11^. To include a fluorescent nuclear stain, propidium iodide (Abcam, 1:100) was added to the LCO staining solution and co-incubated for the entire duration. Subsequently, sections were incubated with TrueBlack Quencher (Biotium, 1:20 in 70% ethanol) for 10 s, according to manufacturer’s instructions, in order to reduce autofluorescence caused by lipofuscin and other tissue components. The sections were then dried at RT for 20 min and coverslipped with fluorescence mounting medium (DAKO), Art. No. S302380-2). The slides were dried overnight at RT and stored at 4°C until imaging.

To obtain an overview of overall tissue integrity and the distribution of ischemic neurons, one neighboring brain section per patient was stained with cresyl violet (Morphisto), according to standard protocols. Confocal imaging was performed with a Stellaris 5 microscope (Leica). All Aβ plaques identified throughout the entire human brain sections (based on hFTAA signal) were imaged and 30 µm Z-stacks were acquired with a 0.5 µm step size. Images were acquired with a HC PL APO CS22 40x/1.3 OIL objective and exported with a bit depth of 8. Using Imaris (Bitplane, V.10.2)., 3D surface reconstructions were created for every channel based on a fixed intensity threshold. Microglia were reconstructed using Iba1 signal, and Aβ plaques were reconstructed based on hFTAA (total Aβ plaque) and qFTAA (Aβ plaque core) signals, respectively. Reconstructed microglia were automatically categorized into “plaque-associated” and “plaque-distant” microglia, based on the smallest distance to an Aβ plaque (< 13 µm or ≧ 13 µm, respectively). Brightfield images were obtained with a SLIDEVIEW VS200 slide scanner (Olympus) at 20x magnification. Imaris data post-processing was performed in R Studio (V. 2024.12.0+467). Slide scanner images were analyzed and exported using OlyVIA software (Olympus, V.3.8).

